# A heterotrimeric complex of *Toxoplasma* proteins promotes parasite survival in interferon gamma stimulated human cells

**DOI:** 10.1101/2022.12.08.519568

**Authors:** Eloise J. Lockyer, Francesca Torelli, Simon Butterworth, Ok-Ryul Song, Steven Howell, Anne Weston, Philip East, Moritz Treeck

**Affiliations:** Signalling in Apicomplexan Parasites Laboratory, The Francis Crick Institute, London, UK; High-Throughput Screening Science Technology Platform, The Francis Crick Institute, London, United Kingdom; Proteomics Science Technology Platform, The Francis Crick Institute, London, United Kingdom; Electron Microscopy Science Technology Platform, The Francis Crick Institute, London, United Kingdom; Bioinformatics and Biostatistics Science Technology Platform, The Francis Crick Institute, London, United Kingdom

## Abstract

*Toxoplasma gondii* secretes protein effectors to subvert the human immune system sufficiently to establish a chronic infection. Relative to murine infections, little is known about which parasite effectors disarm human immune responses. Here we used targeted CRISPR screening to identify secreted protein effectors required for parasite survival in IFNγ-activated human cells. Independent screens were carried out using two *Toxoplasma* strains which differ in virulence in mice, leading to the identification of effectors required for survival in IFNγ-activated human cells. We identify the secreted protein GRA57 and two other proteins, GRA70 and GRA71, that together form a complex which enhances the ability of parasites to persist in IFNγ-activated human foreskin fibroblasts (HFFs). Components of the protein machinery required for export of *Toxoplasma* proteins into the host cell were also found to be important for parasite resistance to IFNγ in human cells, but these export components function independently of the identified protein complex. Host-mediated ubiquitination of the parasite vacuole has previously been associated with increased parasite clearance from human cells, but we find that vacuoles from GRA57, GRA70 and GRA71 knockout strains are surprisingly less ubiquitinated by the host cell. We hypothesise that deletion of this trimeric complex renders parasites hypersensitive to remaining ubiquitination, resulting in increased parasite clearance.

## Introduction

*Toxoplasma gondii* is an obligate intracellular parasite that can infect any nucleated cell of virtually any warm-blooded animal. With an estimated human worldwide seroprevalence of 30%, it is also likely one of the most prevalent human protozoan parasites (Torgerson and Mastroiacovo, 2013). Though *Toxoplasma* can cause disease in immunocompromised patients or following congenital transmission, most immunocompetent individuals control the infection through the combined action of the adaptive and innate immune systems.

The *Toxoplasma gondii* lifecycle comprises two distinct phases; a sexual stage that takes place exclusively in the definitive feline host, and an asexual stage that can occur in a very broad range of intermediate hosts (Elmore *et al*., 2010). The evolutionary importance of an intermediate host is intrinsically linked to the frequency with which it promotes transmission of *Toxoplasma* parasites back to the feline definitive host (Gazzinelli *et al*., 2014). Rodent species and other small warm-blooded animals that are prey to felines are therefore important natural intermediate hosts for *Toxoplasma*, but as humans are very rarely natural feline prey, they are considered ‘accidental’ intermediate hosts.

Once within an intermediate host, *Toxoplasma* actively invades host cells and resides in a host-derived membrane-bound vacuole known as the parasitophorous vacuole (PV). In mice and humans, host cell-autonomous immunity to *Toxoplasma* is mediated by the type II interferon IFNγ. IFNγ stimulation leads to the upregulation of hundreds of IFN-stimulated genes (ISGs) through STAT1-induced transcription of gamma-activated sequence elements (MacMicking, 2012; Malterer, Glass and Newman, 2014). To counteract the IFNγ response, *Toxoplasma* secretes protein effectors from specialised rhoptry and dense granule organelles. Dense granule proteins (GRAs) and rhoptry proteins (ROPs) transform the host cell by altering gene transcription and host metabolism, mediating the acquisition of nutrients, removing host cell factors from the PV membrane (PVM) and targeting pro-inflammatory signalling pathways (Reviewed in (Melo, Jensen and Saeij, 2011; Hunter and Sibley, 2012; Hakimi, Olias and David, 2017; Matta *et al*., 2021; Panas and Boothroyd, 2021; Tomita *et al*., 2021)). Global mapping of the *Toxoplasma* proteome using hyperplexed localisation of organelle proteins by isotope tagging (hyperLOPIT) has predicted the existence of at least 124 GRAs and 106 ROPs (Barylyuk *et al*., 2020). Sequence polymorphisms and differential expression of secreted effectors across the three main clonal lineages of *Toxoplasma* (types I, II & III) have been shown to determine differences in strain virulence (Ajzenberg *et al*., 2004; Swierzy *et al*., 2017), however the vast majority of ROPs and GRAs remain uncharacterised.

GRAs can be translocated across the parasite plasma membrane and PVM into the host cell cytoplasm via the MYR complex, a multi-protein putative translocon. To date, eight proteins have been identified as necessary for protein translocation: the putative MYR translocon components; MYR1, MYR2, MYR3, MYR4 and ROP17 (Franco *et al*., 2016; Marino *et al*., 2018; Panas *et al*., 2019; Cygan *et al*., 2020), as well as GRA44, GRA45 and ASP5 (Coffey *et al*., 2015; Cygan *et al*., 2020). The secretion of MYR-dependent effectors is responsible for the vast majority of host cell transcriptome changes induced by *Toxoplasma* infection in both human and mouse cells (Melo *et al*., 2013; Naor *et al*., 2018). MYR1 knockouts display an expected reduction in virulence *in vivo*, but this phenotype is lost in pooled knockout CRISPR screens, suggesting MYR1-dependent effectors may exert paracrine effects at the site of infection (Young *et al*., 2019). How the MYR proteins directly facilitate protein translocation is not yet understood, and while MYR1 and MYR3 stably associate *in vitro* (Marino *et al*., 2018), it is not yet clear whether all MYRs form a single complex required for protein translocation.

Efforts to characterise the function of *Toxoplasma*-secreted effectors historically centred on their role in the murine host, leading to the discovery of ROP18, ROP5 and ROP17 (Saeij *et al*., 2006; Taylor *et al*., 2006; Behnke *et al*., 2011; Reese *et al*., 2011). In mice, these effectors act cooperatively to prevent the loading of the immunity-related GTPase (IRG) family of large GTPases (Fentress *et al*., 2010; Steinfeldt *et al*., 2010; Behnke *et al*., 2012; Fleckenstein *et al*., 2012; Niedelman *et al*., 2012; Etheridge *et al*., 2014), which are strongly induced by IFNγ in multiple mouse cell types (Khaminets *et al*., 2010; Howard, Hunn and Steinfeldt, 2011). IRG proteins load sequentially onto the parasitophorous vacuole membrane (PVM) and cooperatively oligomerise to vesiculate and rupture the membrane, leading to parasite clearance and necrotic host cell death (Martens *et al*., 2005; Zhao *et al*., 2009; Khaminets *et al*., 2010). The allelic combination of ROP18/ROP5/ROP17 in each parasite strain determines the strain-specific virulence of *Toxoplasma* in mice (Behnke, Dubey and Sibley, 2016). While mice have a dramatically expanded family of 23 IRGs, IRGs are largely absent in humans having been mostly lost in the primate lineage prior to the evolution of monkeys (Bekpen *et al*., 2005). As a result, ROP18/ROP17/ROP5 do not function similarly to resist vacuole clearance in human cells (Niedelman *et al*., 2012).

In humans, the IFNγ-induced responses to *Toxoplasma* are highly cell-type specific. These mechanisms include autophagic clearance (Selleck *et al*., 2015), endo-lysosomal destruction (Clough *et al*., 2016; Mukhopadhyay *et al*., 2020), nutrient deprivation (Pfefferkorn, 1984; Dimier and Bout, 1997), or induction of host cell death (Niedelman *et al*., 2013). In human macrophages, the p65 GTPase guanylate binding protein 1 (GBP1) has been demonstrated to mediate killing of *Toxoplasma* through recruitment to and disruption of the PVM, leading to host cell apoptosis (Fisch *et al*., 2019).

In multiple mouse and human cell types, IFNγ-driven ubiquitination of the PVM by host E3 ligases serves as an initial marker for eventual parasite clearance. In mouse embryonic fibroblasts (MEFs) and bone marrow-derived macrophages (BMDMs), TRAF6-mediated ubiquitination of type II and III vacuoles enhances the recruitment of GBPs, resulting in rupture of the PVM (Haldar *et al*., 2015; Mukhopadhyay *et al*., 2020). The E3 ligase TRIM21 has also been shown to promote clearance of type II and III strains *in vivo* (Foltz *et al*., 2017), and ubiquitinates type III vacuoles in human foreskin fibroblasts (HFFs) resulting in parasite growth restriction (Yao *et al*., 2021). In human umbilical vein endothelial cells (HUVECs), K63-linked ubiquitination initiates a cascade of autophagy marker recruitment that culminates in acidification of the parasite vacuole (Clough *et al*., 2016), while in HeLa cells ubiquitination instead leads to recruitment of LC3 and stunting of parasite growth (Selleck *et al*., 2015). Compared to type II & type III parasites, type I parasites are more resistant to ubiquitination and clearance in both HUVECs and HeLa cells. Recent evidence suggests a major function for the E3 ligase RNF213 in ubiquitination of the PVM and enhancing parasite clearance, in multiple human cell types and in a strain-independent manner (Hernandez *et al*., 2022; Matta *et al*., 2022). The importance of each E3 ligase and ubiquitin linkage therefore depends on the exact combination of host species, parasite strain and cell type studied, but nevertheless ubiquitination has emerged as a key process in the regulation of *Toxoplasma* clearance. The *Toxoplasma*-derived targets that are recognised by these host E3 ligases remain to be determined.

*Toxoplasma* effectors that have been described to counteract IFNγ in human cells include the host transcriptional repressor IST, which blocks IFNγ-stimulated gene transcription (Gay *et al*., 2016; Olias *et al*., 2016). However, IST can only prevent clearance in cells that have not been pre-stimulated with IFNγ, a condition likely not met during acute infection. Another effector is NSM, which blocks host cell necroptosis during the bradyzoite stages of infection (Rosenberg and Sibley, 2021). More recently, we have shown that ROP1 counteracts IFNγ immune responses in both murine and human macrophages (Butterworth *et al*., 2022). Finally, in human THP-1-derived macrophages, deletion of the chaperone protein GRA45 increases the sensitivity of parasites to IFNγ-mediated growth inhibition (Y. Wang *et al*., 2020). There therefore remains a significant gap in our understanding of which effectors mediate *Toxoplasma* virulence in humans (Gazzinelli *et al*., 2014; Fisch, Clough and Frickel, 2019).

To address this gap, we performed targeted CRISPR screening of the *Toxoplasma* ‘secretome’ (Barylyuk *et al*., 2020) during infection of unstimulated and IFNγ-stimulated HFFs, using our previously described CRISPR platform (Young *et al*., 2019). Independent experiments were performed in type I (RH) and type II (PRU) parasite strains, which allowed identification of both strain-dependent and independent effectors. We found that GRA57 is a strain-independent effector that protects *Toxoplasma* from IFNγ-mediated vacuole clearance in HFFs. GRA57 was found to interact with two other dense granule proteins, GRA70 (TGME49_249990) and GRA71 (TGME49_309600), which resist IFNγ-mediated vacuole clearance to a similar degree as GRA57, indicating that the three proteins function in a complex. Deletion of any member of this trio results in reduced PVM ubiquitination in HFFs. We also found two components of the MYR translocon, MYR1 and MYR3, contribute to IFNγ resistance in HFFs, though we expect that one or several unidentified MYR-dependent effectors play an additional significant role. As GRA57 does not impact GRA export, and MYR1 and MYR3 do not display a ubiquitination phenotype, we conclude that GRA57 and the MYR proteins function independently of each other. These data suggest that a novel MYR-independent trimeric complex of dense granule proteins localised within the PV contribute to resisting IFNγ-induced vacuole clearance in HFFs.

## Results

### CRISPR screening the *Toxoplasma* secretome in IFNγ-activated human cells

To screen for effector proteins that promote *Toxoplasma* survival in IFNγ-activated human cells, we used a previously established *in vivo* CRISPR screening platform (Young *et al*., 2019) with a guide library targeting 253 predicted or validated dense granule and rhoptry proteins. To allow for identification of strain specific and independent effectors, we generated type I and type II CRISPR knockout parasite pools by transfecting this library into RHΔHXGPRT and PRUΔHXGPRT strains. Transfected parasites were selected in HFFs for 8 days to generate a pooled mutant parasite population. Pooled mutant parasites were then used to infect HFFs that were either untreated or pre-stimulated with IFNγ, for two lytic cycles (96h) (Figure 1A).

**Figure 1:**
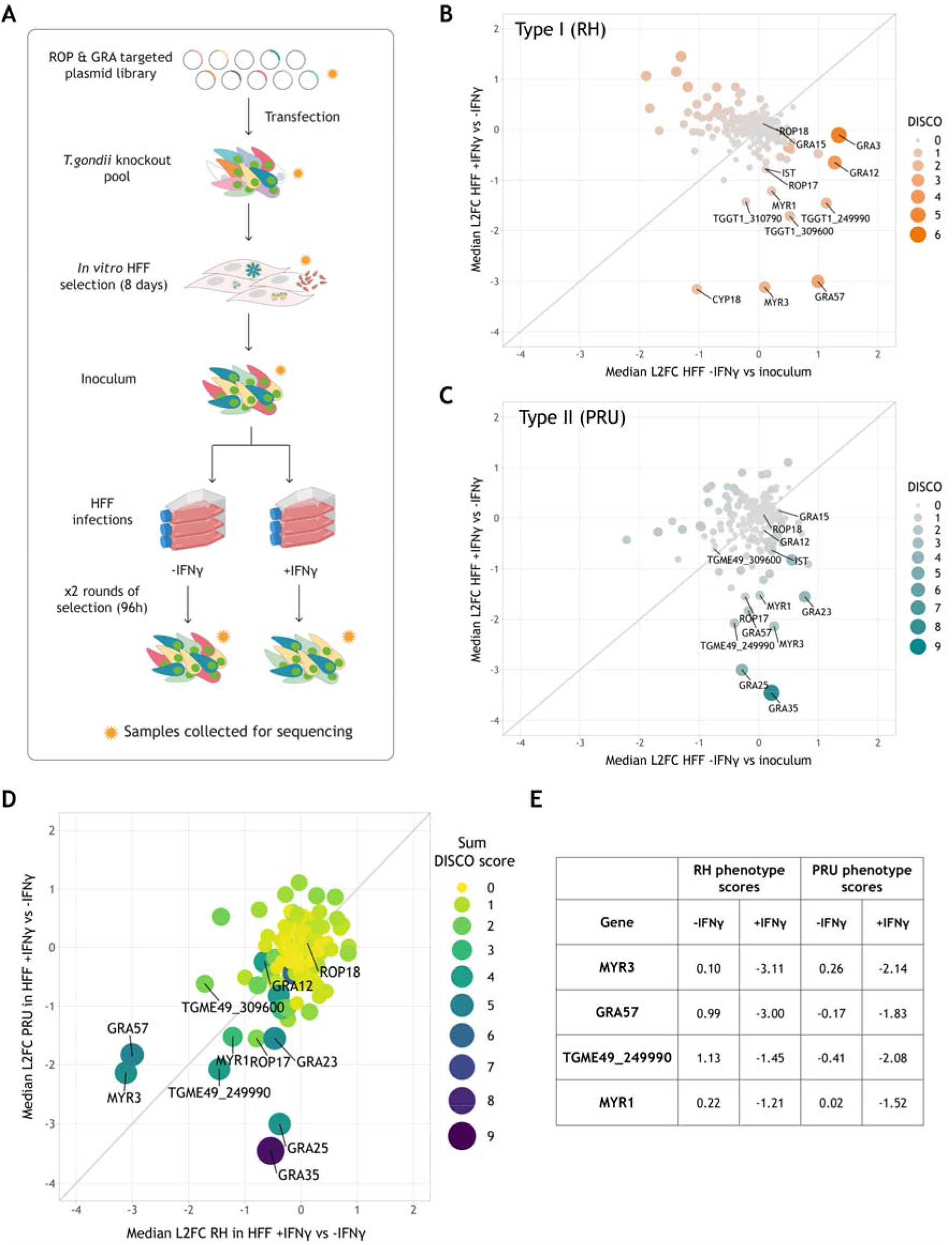
CRISPR screening the *Toxoplasma* secretome to identify novel effectors in IFNγ-activated HFFs. **(A)** Schematic of experimental design. A pCas9-T2A-HXGPRT-sgRNA vector library designed against predicted secreted effectors was transfected into RH or PRUΔHXGPRT to generate a mutant parasite pool. Transfectants were grown under M/X selection in HFFs for 8 days, then combined to generate the inoculum. HFFs were infected in triplicate, in the presence or absence of 24h pre-stimulation with 100U/ml IFNγ. After 48h infection, a subset of parasites from each condition were retrieved and used for a second round of infection for a further 48h. Subsets of parasite pools were taken for gDNA extraction and sequencing at each stage (indicated by orange stars) to determine the relative abundance of guides. **(B & C)** Discordance plots showing median log2 fold changes (L2FCs) for *Toxoplasma* effector genes in **(B)** RH background and **(C)** PRU background. L2FCs for growth in IFNγ-activated HFFs are plotted against control L2FCs for growth in unstimulated cells. Colour and size of data points indicate strength of discordance score. **D)** Correlation between RH and PRU screens in IFNγ-activated HFFs. Genes with L2FCs <-1 for growth in unstimulated HFFs in either screen are excluded. **E)** Table of genes with L2FCs <-1 for growth in IFNγ-activated HFFs in both RH and PRU backgrounds.

To identify genes that confer a growth benefit for *Toxoplasma* in HFFs in the presence of IFNγ, we measured the relative abundance of guide RNAs (gRNAs) targeting each gene in the *Toxoplasma* mutant library before and after selection in IFNγ-treated or untreated HFFs. Sequencing read counts were used to calculate the median Log2 fold change (L2FC) of guides targeting each gene between conditions, and matched gene concordance/discordance (DISCO) to compare gene phenotypes between treatment conditions. To assess which effectors contributed predominantly to survival of IFNγ responses in HFFs, we calculated the L2FC for each gene between IFNγ-stimulated and untreated cells after two rounds of selection, then compared this to *in vitro* growth phenotypes using the L2FC between untreated cells and the starting parasite inoculum.

Parasites were highly restricted in HFFs, with the RH and PRU mutant populations restricted by 85% and 90% respectively after 48h of infection. Results from both the RH and PRU screens showed that ROP18 and GRA12-two major virulence factors in murine cells and *in vivo* (Saeij *et al*., 2006; Niedelman *et al*., 2012; Fox *et al*., 2019; J. Wang *et al*., 2020)-have no major impact on parasite survival in HFFs (Figure 1B & 1C). In the RH screen, effectors with the highest IFNγ phenotype were CYP18, a protein with a proinflammatory effect in macrophages (Ibrahim *et al*., 2009), MYR3, a component of the dense granule export translocon (Marino *et al*., 2018), GRA57, a cyst-localising protein (Nadipuram *et al*., 2020) and TGME49_309600, a hypothetical dense granule protein (Figure 1B). In the PRU screen, the highest scoring effectors were GRA35, an inducer of pyroptosis in rat macrophages (Wang *et al*., 2019), GRA25, an essential virulence factor in mice (Shastri *et al*., 2014), MYR3, TGME49_249990, a predicted dense granule protein and GRA57 (Figure 1C).

To identify strain-independent effectors contributing specifically to survival of HFF IFNγ responses, we compared genes with strong IFNγ phenotype scores in the RH and PRU screens, while excluding those with strong phenotype scores in unstimulated HFFs (Figure 1D & 1E & Supplementary Data 1). For both screens, three components of the MYR translocon-MYR1, MYR3 and ROP17-were among the genes with the highest phenotype scores for survival in IFNγ-activated HFFs. As the export of many dense granule proteins into the host cell has been shown to be MYR-dependent, it is likely that the role of MYR components in IFNγ resistance results from the pleiotropic effect of abrogating dense granule export. We instead focused our efforts on the further characterisation of GRA57, which showed the second strongest phenotype in both screens and had no previously known function.

### GRA57 is an intravacuolar protein required for *Toxoplasma* survival of IFNγ responses in HFFs

To investigate the function of GRA57, we generated a knockout line in the type I RH strain via Cas9-mediated gene disruption and integration of an mCherry-T2A-HXGPRT repair cassette. We complemented the GRA57 coding sequence with a C-terminal HA tag back into this line at the non-essential *uprt* locus (Figure S1A & S1B). This allowed us to confirm the localisation of GRA57-HA described in (Nadipuram *et al*., 2020) via immunofluorescence, showing that GRA57 resides in the intravacuolar network (IVN) (Figure 2A), and partially co-localises with GRA3 (Figure S3), a marker of the PVM (Deffieu *et al*., 2019). GRA57 is a relatively large dense granule protein predicted to be 246kDa, which we confirmed via western blot (Figure 2B).

**Figure 2:**
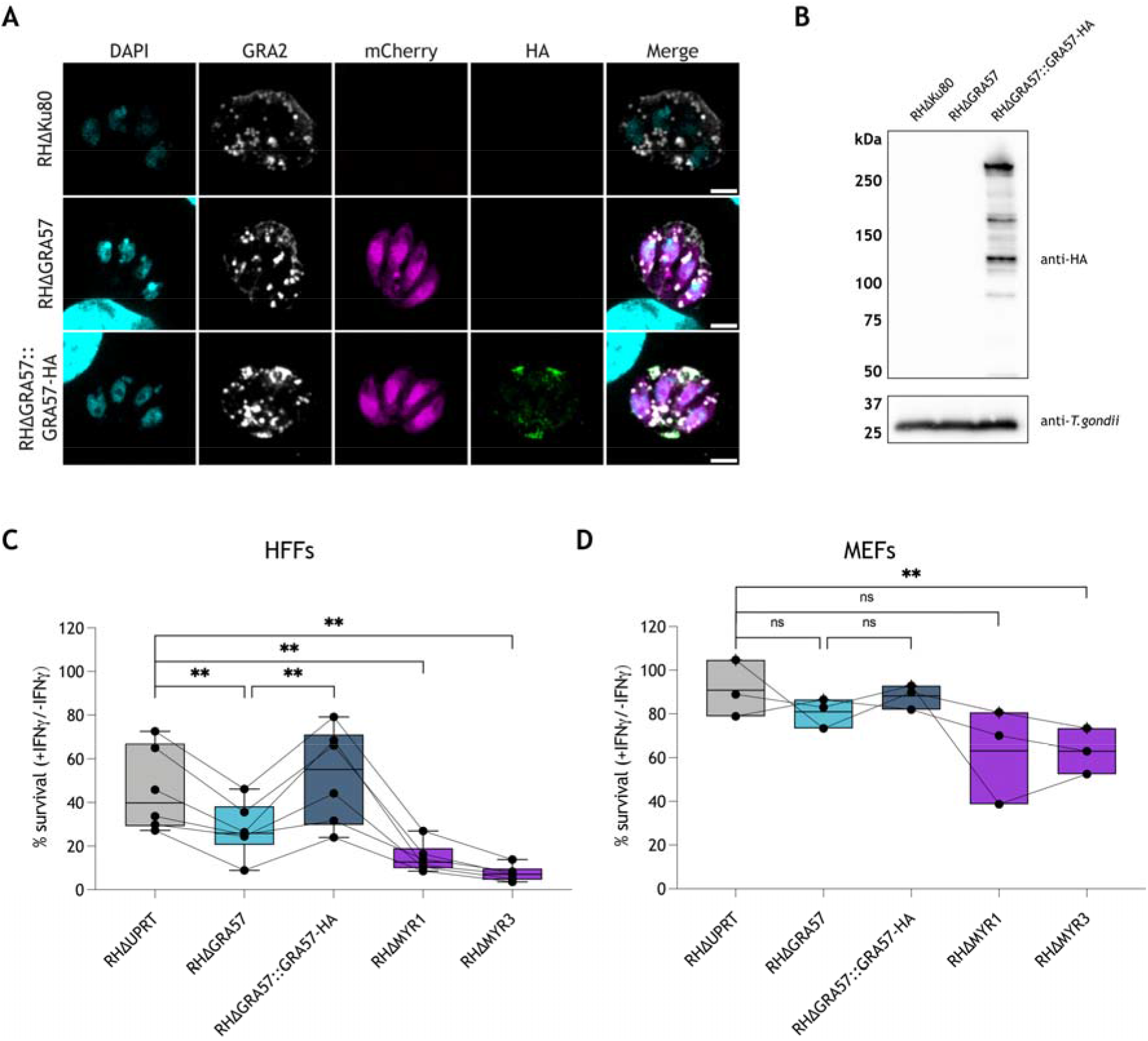
GRA57 is an intravacuolar protein that contributes to parasite survival of IFNγ in HFFs. **(A)** IFA of RHΔGRA57 and RHΔGRA57::GRA57-HA lines generated for this study. HA-tagged GRA57 co-localises with GRA2-a marker of the IVN. Scale bar= 3μm. **(B)** Western blot analysis of GRA57-HA shows it is a 250kDa protein that is relatively highly expressed **(C & D)** Live restriction assays in **(C)** HFFs and **(D)** MEFs using mCherry fluorescence area as a readout for parasite survival. Host cells were pre-stimulated for 24h with 100U/ml IFNγ or left untreated then infected in technical triplicate with the indicated parasite strains for 48h (HFFs) or 24h (MEFs) at an MOI of 0.3. Infected cells were then imaged live on a Cytation plate reader. Total mCherry signal area per well was measured to determine parasite growth in IFNγ stimulated relative to unstimulated cells. Data displayed as median survival with individual biological replicates overlayed. p-values were calculated by paired two-sided t-test. **, p < 0.01; ns, not significant.

To verify the phenotype observed in the CRISPR screens, we assessed the survival of the RHΔGRA57 and RHΔGRA57::GRA57-HA strains in IFNγ-activated HFFs. HFFs were pre-stimulated with 100U/ml IFNγ for 24h prior to infection with the mCherry expressing parasite lines, or left untreated as a control. As a wildtype control, we used a parental strain where an mCherry-T2A-HXGPRT cassette was introduced into the *uprt* locus. The total mCherry signal area was measured from microscopy images taken on a Cytation plate imager as a readout for *Toxoplasma* survival, and the percentage survival was calculated as a proportion of the area in unstimulated vs IFNγ-stimulated cells. We also generated a RHΔMYR3 strain and included this alongside RHΔMYR1 given the strong phenotype observed for these effectors in the pooled CRISPR screens.

In IFNγ-stimulated HFFs, RHΔGRA57 parasites displayed a relative decrease in survival of 39% compared to RHΔUPRT controls (Figure 2C). This decrease in survival was rescued in the complemented line, confirming that the absence of GRA57 renders parasites more susceptible to IFNγ restriction in HFFs. RHΔMYR3 and RHΔMYR1 were more highly restricted, with relative reductions in survival of 68% and 84%, respectively. This indicates that while GRA57 plays an important role in resisting host cell restriction, it is unlikely to have a major influence on MYR-dependent protein export. This is further investigated later in this manuscript.

Since GRA57 has not previously been identified as an important effector for survival in mice (Sangaré *et al*., 2019; Y. Wang *et al*., 2020), this indicates that GRA57 plays a role in only some species, or specific cell types. To verify this, we performed restriction assays in IFNγ-activated MEFs as an equivalent murine cell type. Survival levels for all parasite lines were higher in MEFs (91% for RHΔUPRT) than in HFFs (46% for RHΔUPRT), suggesting that HFFs restrict type I parasites more effectively than MEFs, at least under the conditions tested (Figure 2D). This is in line with previously published data that shows type I parasites are highly resistant to restriction in MEFs (Niedelman *et al*., 2013).

RHΔGRA57 parasites displayed a relative decrease in survival of 11%, however this was not a significant change, indicating that GRA57 is not as important for survival in MEFs as in HFFs. In contrast, RHΔMYR3 parasites had a significant relative reduction in survival of 31%, with RHΔMYR1 showing a similar, but not statistically significant trend of 30% relative reduction, indicating that preventing effector export has a major impact on parasite survival regardless of host species. Overall, this data verifies that GRA57 and components of the MYR translocon are novel *Toxoplasma* survival factors that contribute to parasite resistance to IFNγ in HFFs.

### GRA57 forms a complex with TGGT1_249990 and TGGT1_309600

To identify interaction partners of GRA57 during infection of HFFs, which could indicate how GRA57 functions to protect the parasite, we performed coimmunoprecipitation (IP) experiments with an endogenously tagged RHGRA57-HA strain (Figure S2). To enable detection of protein-protein interactions specific to activated cells, HFFs were pre-stimulated before infection with 2.5U/ml IFNγ for 6h, which we found was sufficient to induce IFNγ responses while retaining host cell viability. HFFs were infected for 24h with either RHΔKu80 or RHGRA57-HA parasites prior to lysis. GRA57-HA was immunoprecipitated from lysates, then coimmunoprecipitated proteins were identified by liquid chromatography (LC)-tandem mass spectrometry (MS/MS).

GRA57 was significantly enriched in RHGRA57-HA infected samples relative to RHΔKu80 infected samples (Figure 3A). Strikingly, the two most strongly coenriched proteins in the tagged samples were TGGT1_249990 and TGGT1_309600 (Figure 3A)-two predicted dense granule proteins that also displayed IFNγ-specific phenotypes in both CRISPR screens (Figure 1D and Supplementary Data 1). The mean label-free quantification (LFQ) values detected for GRA57, TGGT1_249990 and TGGT1_309600 were similar (Figure 3B), which strongly suggests that GRA57, TGGT1_249990, and TGGT1_309600 form a heterotrimeric complex. Because both TGGT1_249990 and TGGT1_309600 are predicted dense granule proteins that interact with GRA57, they are referred to as GRA70 and GRA71, respectively, from here onwards. All three proteins are predicted to have transmembrane helices, with GRA57 and GRA70 also showing regions of predicted coiled-coil towards the N-terminus (Figure 3C).

**Figure 3:**
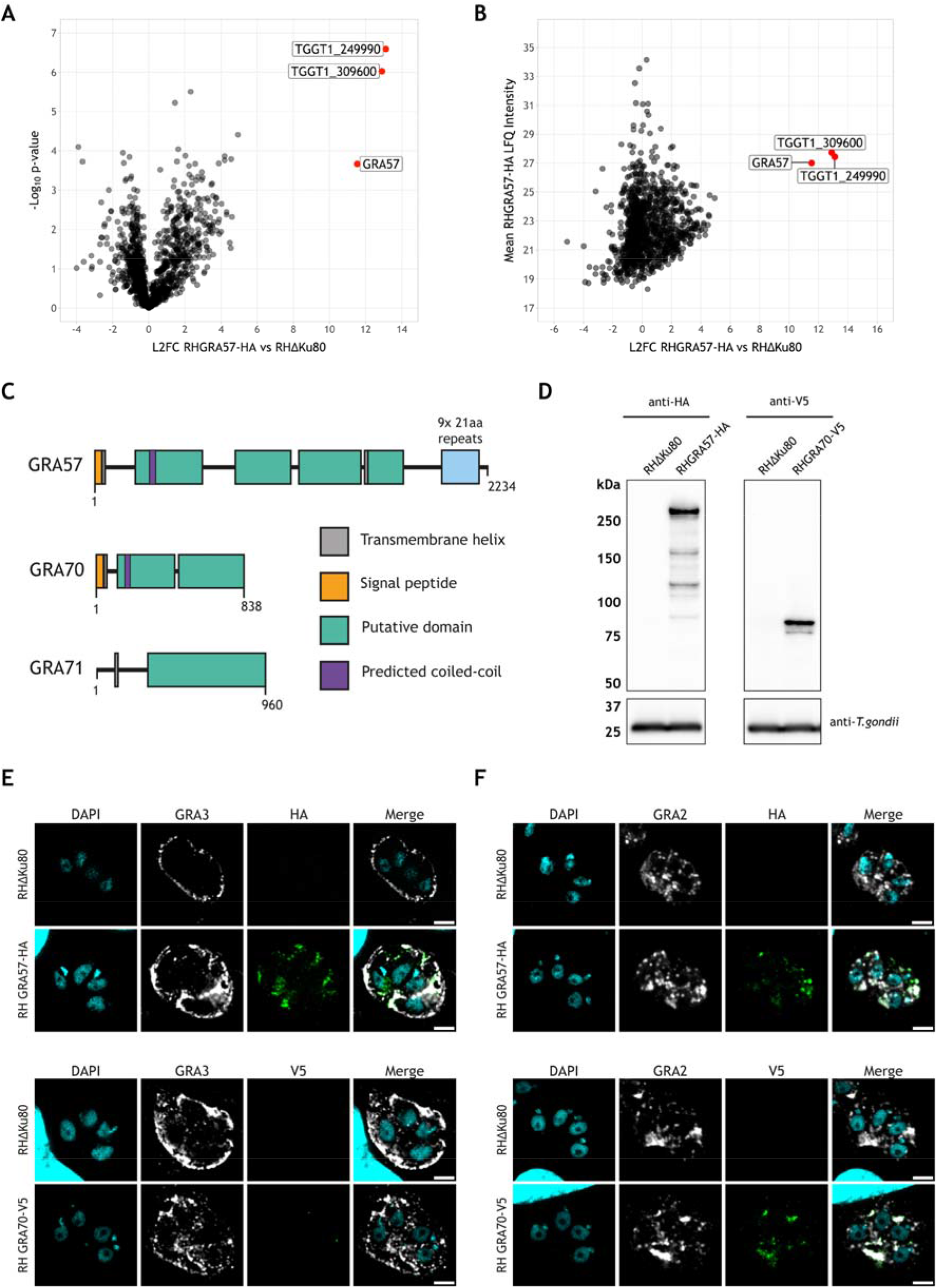
CoIP of GRA57-HA shows it likely forms a complex with TGGT1_249990 and TGGT1_309600 within the PV. **(A,B)** Co-immunoprecipitation of endogenously HA-tagged RHGRA57. HFFs were pre-stimulated with 2.5U/ml IFNγ for 6h, then infected in triplicate with either RHΔKu80 or RHGRA57-HA for a further 24h. GRA57 was immunoprecipitated with anti-HA agarose matrix then immunoprecipitate was analysed by mass spectrometry. **(A)** Enrichment of proteins in RHGRA57-HA vs RHΔKu80 control samples. **(B)** Mean LFQ signal intensity detected for proteins in RHGRA57-HA samples. **(C)** Schematic of putative domains in GRA57, GRA70 and GRA71, to scale. Domains and boundaries were assigned based on regions of predicted secondary structure using PsiPred. Transmembrane helices are annotated where predicted by at least two programs (MEMSAT-SVM, DeepTMHMM and CCTOP). Regions of predicted coiled-coil were determined using Waggawagga. **(D)** Western blot protein expression analysis of endogenously tagged lines. **(E)** Co-localisation of endogenously c-terminally tagged GRA57 and GRA70 with the PVM marker GRA3. Samples were permeabilised with 0.1% saponin. Scale bar= 3μm. **(F)** Co-localisation of endogenously c-terminally tagged GRA57 and GRA70 with the IVN marker GRA2. Samples were permeabilised with 0.2% Triton-X100. Scale bar= 3μm.

To show further evidence of this complex, we determined the localisation of GRA70 via immunofluorescence. GRA70 was C-terminally tagged with a V5 epitope to validate its localisation in the PV (Figure S2). Western blotting showed the expected size of GRA70 (93kDa) (Figure 3D), and immunofluorescence confirmed that GRA70 is secreted into the parasitophorous vacuole (Figure 3E & 3F). GRA70 and GRA57 display similar patterns of localisation in the IVN at 24h post-infection, partially co-localising with the IVN marker GRA2 (Sibley *et al*., 1995) (Figure 3F). With saponin permeabilisation we could only detect minor signal for GRA70-V5 in the vacuole (Figure 3E), in contrast to GRA57-HA which is detectable with both saponin and triton permeabilisation (Figure 3E & 3F). This suggests the C-terminal portion of GRA57 is exposed to the host cell cytosol, whereas the GRA70 C-terminus remains within the PV. We did not detect any signal beyond the PVM for either GRA70 or GRA57, suggesting that these effectors are not secreted into the host cell. Together this data suggests that GRA57, GRA70 and GRA71 form a complex that is anchored in the PVM, or IVN, via GRA57.

### GRA57, GRA70 and GRA71 resist vacuole clearance in IFNγ-activated HFFs

To verify the phenotype seen in CRISPR screens for GRA70 and GRA71, we generated individual knockouts of GRA71 and GRA70 in the RH strain (Figure S1). To assess with higher precision whether these parasites are more susceptible to IFNγ-induced growth restriction or vacuole clearance, we analysed the survival of these lines in IFNγ-activated HFFs using a high content imaging method previously established in (Butterworth *et al*., 2022).

GRA57 knockouts displayed increased sensitivity to IFNγ in HFFs (Figure 4A & 4B), concurrent with the phenotype observed in the CRISPR screen (Figure 1B). RHΔGRA70 and RHΔGRA71 parasites were restricted to the same level as RHΔGRA57 parasites (mean relative reduction in overall parasite numbers vs RHΔUPRT-30%, 33% and 31%, respectively) confirming that these are also resistance factors in activated HFFs (Figure 4A). For GRA57, GRA70 and GRA71, the overall reduction in parasite survival could mainly be attributed to a reduction in the number of parasite vacuoles (Figure 4B), with no significant decrease in the size of parasite vacuoles (Figure 4C). This suggests these knockout lines are more susceptible to active clearance, early egress from the host cell (Niedelman *et al*., 2013), or increased host cell death, rather than restriction of parasite growth. For the MYR component knockouts, parasites numbers were reduced as a result of a decrease in both vacuole number and size (Figure 4B & 4C), which may be expected given the combinatorial effects of preventing secretion of effectors that counteract both host pathways. Given that deletion of all three components of the complex leads to a very similar reduction in resistance to vacuole clearance (mean relative reduction in vacuole number vs RHΔUPRT-GRA57: 33%, GRA70: 34%, GRA71: 41%), we conclude that GRA57, GRA70 and GRA71 form a complex that mediates resistance to vacuole clearance in IFNγ-stimulated HFFs.

**Figure 4:**
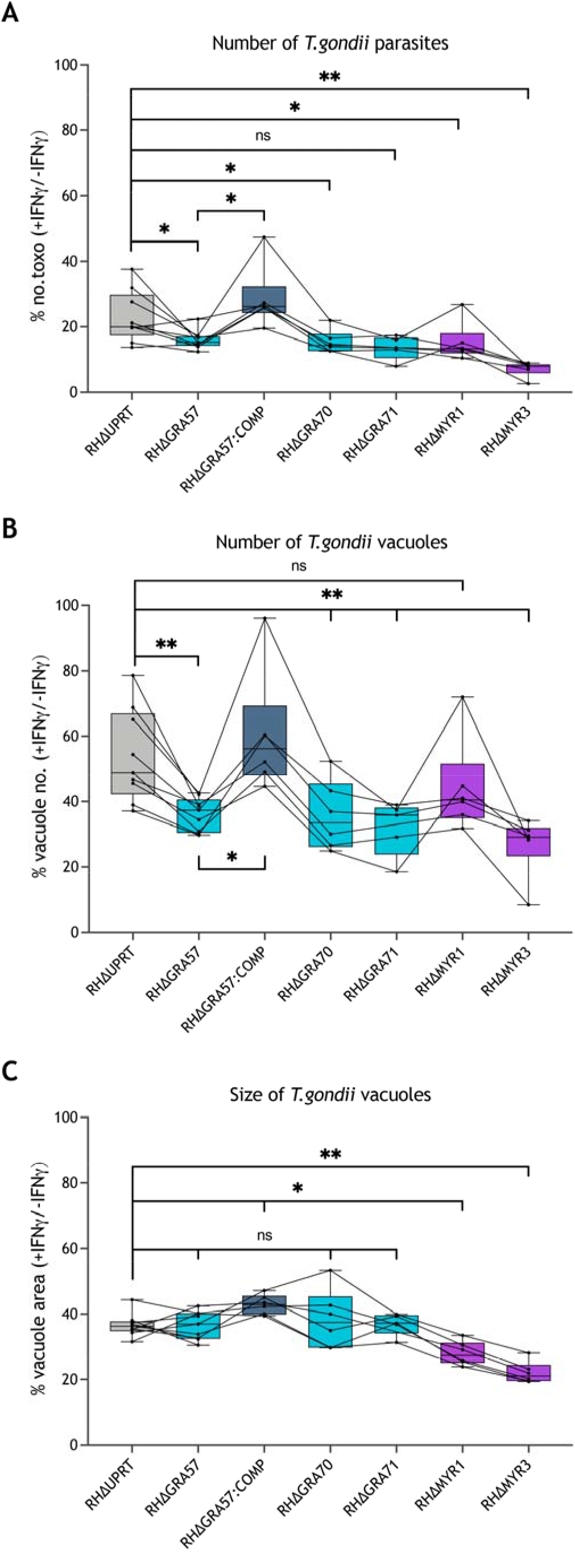
GRA57/GRA70/GRA71 all contribute equally to resisting IFNγ-induced vacuole destruction in human fibroblasts. **(A-C)** HFFs cells were pre-stimulated for 24h with 100U/ml IFNγ or left untreated then infected in technical triplicate with the indicated parasite strains for 24h at an MOI of 0.3. Parasite survival was quantified through automated high content imaging. (**A)** Parasite survival calculated as total number of *Toxoplasma* in IFNγ-stimulated cells as a percentage of the total in unstimulated cells. (**B)** Parasite survival calculated as number of vacuoles in IFNγ-stimulated cells as a percentage of the number in unstimulated cells. **(C)** Parasite survival calculated as mean vacuole area per well [px^2^] in IFNγ-stimulated cells as a percentage of the area in unstimulated cells. Data displayed as median survival with individual biological replicates overlayed. p-values were calculated by paired two-sided t-test. *, p < 0.05; **, p < 0.01; ns, not significant.

### GRA57 does not affect tryptophan metabolism, effector export or host cell gene expression

To determine the mechanism through which GRA57 protects parasites from IFNγ defences, we investigated the role of tryptophan metabolism in the increased susceptibility of RHΔGRA57 parasites in HFFs. Depletion of tryptophan, for which *Toxoplasma* is auxotrophic, has previously been described as an IFNγ-induced restriction mechanism in HFFs. IFNγ-induced expression of the enzyme indoleamine 2,3-dioxygenase (IDO1) results in degradation of intracellular tryptophan, limiting the growth of *Toxoplasma* parasites (Pfefferkorn, 1984; Pfefferkorn, Eckel and Rebhun, 1986; Bando *et al*., 2018). To test the role of GRA57 in tryptophan-dependent parasite restriction we supplemented the HFF growth media with l-tryptophan (L-Trp) as previously described in (Niedelman *et al*., 2013). No rescue of the increased IFNγ susceptibility of RHΔGRA57 parasites was observed, suggesting GRA57 does not act through this pathway (Figure S4). We observed no major rescue in parasite survival for the RHΔUPRT strain upon L-Trp supplementation, indicating that tryptophan metabolism is not the primary restriction mechanism in the HFFs used here. This is in line with other studies that have shown this pathway plays a minimal role in restriction of parasite growth in some HFFs, a discrepancy potentially dependent on the origin of the cells used (Niedelman *et al*., 2013; Mukhopadhyay *et al*., 2020).

We next assessed if GRA57 played a role in dense granule effector export into the host cell. Using RHΔMYR1 and RHΔMYR3 parasites as positive controls, we measured the ability of RHΔGRA57 parasites to induce host c-Myc nuclear translocation via IFA, which in wildtype parasites is induced through MYR-dependent export of GRA16 to the host nucleus (Panas and Boothroyd, 2020). Unlike the near ablation of c-Myc nuclear signal observed upon infection with RHΔMYR1 or RHΔMYR3 parasites, as previously published (Franco *et al*., 2016; Marino *et al*., 2018), at 24h post-infection, there was no significant reduction in c-Myc nuclear signal with RHΔGRA57 parasites (Figure S5). As GRA57 and GRA70 localise to the IVN and PVM, we next assessed the ultrastructure of RHΔGRA57 parasites via transmission electron microscopy (TEM), which showed that the IVN and PV form normally in this strain (Figure S6). Together this confirms that GRA57 is not directly important for MYR dependent protein translocation or biogenesis of the parasite vacuole.

To comprehensively determine if GRA57 has any effect, directly or indirectly, on the host cell transcriptional response to *Toxoplasma*, we compared transcript levels using RNA-Seq of infected HFFs that were untreated or pre-treated with IFNγ. While strong differential gene expression was observed in HFFs upon IFNγ treatment, very few genes were differentially expressed upon infection with RHΔGRA57 parasites compared to WT parasites (Supplementary Data 3). Of the genes that were differentially expressed, expression was not restored to WT infection levels upon infection with RHΔGRA57::GRA57-HA parasites. Collectively this shows that GRA57 plays no role in protein export and does not function through altering host gene expression.

### Deletion of GRA57/GRA70/GRA71 reduces ubiquitination on the parasite vacuole

Ubiquitination of the PVM is a marker for parasite clearance in IFNγ-activated human cells (Clough *et al*., 2016; Yao *et al*., 2021; Hernandez *et al*., 2022; Matta *et al*., 2022). We therefore assessed if deletion of GRA57, GRA70 or GRA71 leads to differences in ubiquitination of the PVM. To test this, we measured total ubiquitin recruitment at 3h post infection. Recruitment was automatically scored by measuring the ubiquitin signal around each vacuole relative to the cytoplasmic background (Figure 5A).

**Figure 5:**
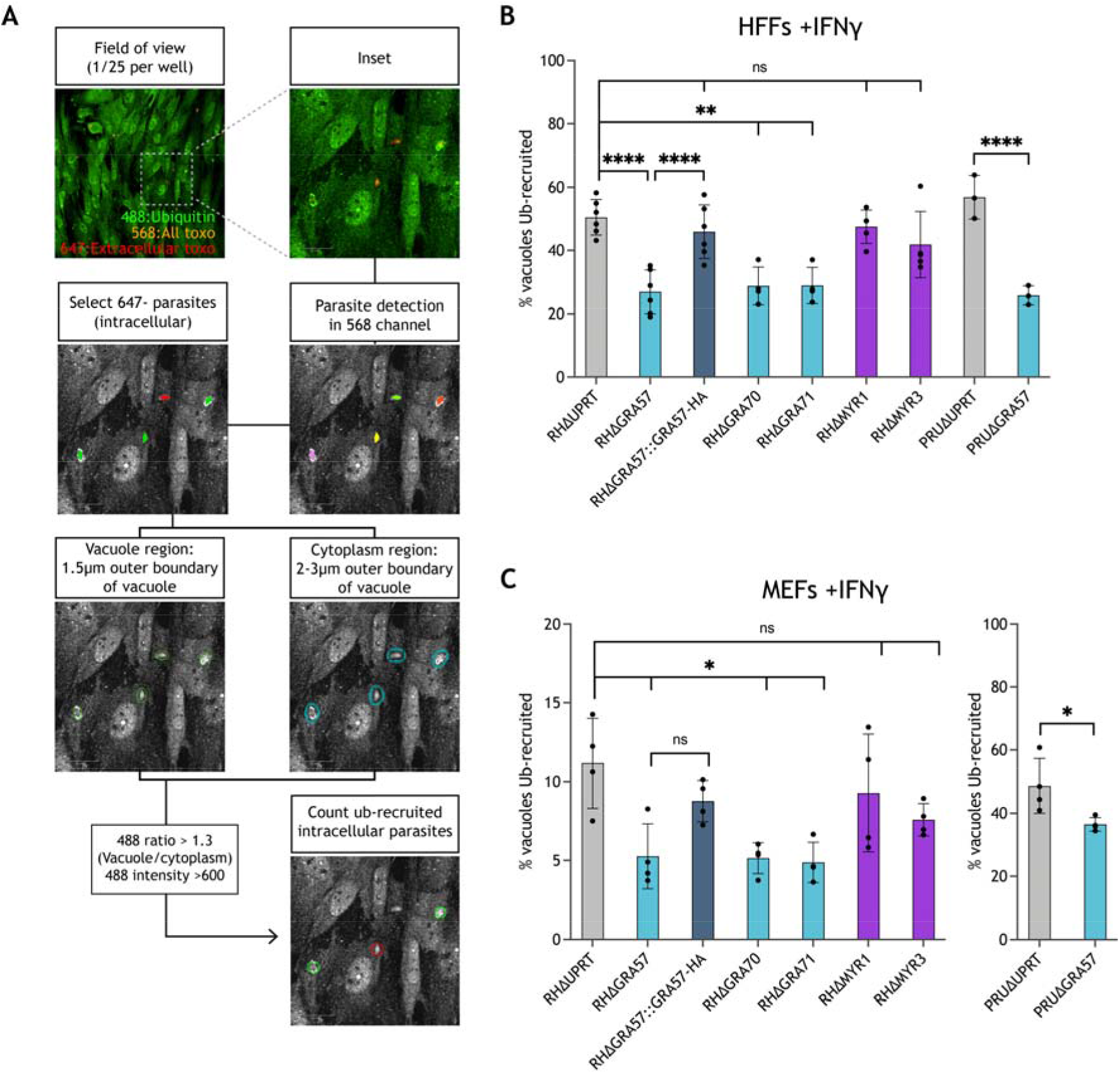
Deletion of GRA57, GRA70 or GRA71 leads to reduced host ubiquitin recruitment to the PVM. **(A)** Schematic of automated high-content imaging analysis pipeline to determine ubiquitin recruitment levels. **(B,C)** Recruitment of host ubiquitin to *Toxoplasma* vacuoles in (**B)** HFFs and (**C)** MEFs. Host cells were pre-stimulated with 100U/ml IFNγ for 24h, infected with indicated lines for 3h prior to fixation and staining for total ubiquitin. Recruitment of ubiquitin was automatically counted using high content imaging and analysis. p-values were calculated by paired two-sided t-test. *, p < 0.05; **, p < 0.01; ***, p < 0.001; ****, p < 0.0001; ns, not significant.

Surprisingly, we found that cells infected with knockouts of GRA57 or its two interaction partners GRA70 and GRA71 had a significantly lower proportion of ubiquitin-recruited vacuoles in both unstimulated (Figure S7A) and IFNγ-pre-stimulated HFFs (Figure 5B). This reduction was also present for PRUΔGRA57 relative to PRUΔUPRT, indicating that this is a strain-independent effect. For RHΔGRA57, the reduction in ubiquitin targeting was restored to wildtype levels upon complementation. In contrast, RHΔMYR1 and RHΔMYR3 vacuoles were equally susceptible to ubiquitin recruitment as RHΔUPRT, providing further support that the GRA57 containing protein complex functions independently of the MYR translocon and its exported effectors.

Given that we observed only a minor increase in restriction in murine fibroblasts upon GRA57 deletion, we next investigated the ubiquitin recruitment levels to these strains in MEFs. Recruitment levels were much higher in PRU than RH, a phenotype which has previously been shown to be dependent on the strain specific expression of GRA15 (Mukhopadhyay *et al*., 2020). Despite overall lower levels of ubiquitin recruitment, deletion of GRA57, GRA70 or GRA71 in the RH strain led to a similar relative reduction in ubiquitin-targeting in IFNγ-activated MEFs as in HFFs of ~50% (Figure 5C). This reduction was again rescued for RHΔGRA57 upon complementation. PRUΔGRA57 displayed a smaller but still significant reduction in ubiquitination relative to wild type in comparison to HFFs. In unstimulated MEFs, the percentage of ubiquitin-positive vacuoles was lower than in unstimulated HFFs, however deletion of GRA57, GRA70 or GRA71 still resulted in a trend towards reduced ubiquitin recruitment (Figure S7B). Together this data suggests that as RHΔGRA57 parasites are much more sensitive to IFNγ-mediated clearance in HFFs than in MEFs, this decrease in ubiquitin recruitment does not correlate with increased susceptibility to IFNγ-induced clearance and may therefore not be directly linked to the parasite clearance mechanism.

Recent work has shown that parasites are cleared in HFFs through endolysosomal acidification of the PV and subsequent killing of the parasites (Mukhopadhyay *et al*., 2020). We hypothesised that RHΔGRA57 vacuoles could be initially ubiquitinated to the same level as wild-type parasites, but cleared by the host cell prior to fixation and staining at 3h. To exclude this possibility, we performed the same recruitment analysis in HFFs at 1 and 2 hours post-infection to determine if ubiquitin recruitment to RHΔGRA57 vacuoles peaked at an earlier timepoint. This showed that RHΔGRA57 vacuoles are consistently less targeted by ubiquitin than RHΔUPRT at these early stages post-infection, in both the untreated and prestimulated HFFs (Figure S8). Together this data confirmed that RHΔGRA57 vacuoles are continuously less susceptible to ubiquitin targeting following host cell invasion.

## Discussion

In this study, we used targeted CRISPR screens to identify *Toxoplasma*-secreted virulence factors that protect the parasite from human cell-autonomous immunity. CRISPR screens were carried out in RH and PRU *Toxoplasma* strains, revealing a novel complex of three interacting dense granule effectors, comprising GRA57, GRA70 & GRA71, that contribute equally to *Toxoplasma* resistance to IFNγ-induced responses in HFFs.

In addition to GRA57 and its two partner proteins, we show that two components of the MYR effector export machinery, MYR1 and MYR3, are important for parasite survival in IFNγ-activated HFFs in both pooled and single knockout infections. Other MYR components including the kinase ROP17 (Panas *et al*., 2019) displayed an IFNγ-survival phenotype in both screens, whereas the effector chaperone GRA45 (Y. Wang *et al*., 2020), was essential for IFNγ survival in only the PRU screen. Together this would suggest that the GRA protein translocation machinery of *Toxoplasma* is required for survival in IFNγ-treated HFFs. However, MYR2 and MYR4 displayed no phenotype in either of our screens, though it is possible these are false negative results. Only MYR1 and MYR3 have previously been shown to stably associate with each other *in vitro* (Marino *et al*., 2018), therefore whether MYR1 and MYR3 have an important additional function in HFFs that is distinct from effector export would be interesting to explore in the future.

In contrast, neither ROP18 nor GRA12 had a major phenotype in IFNγ-stimulated HFFs, whereas in murine screens ROP18 and GRA12 consistently emerge as the most important secreted effectors for parasite survival (Young *et al*., 2019; Y. Wang *et al*., 2020; Butterworth *et al*., 2022). This emphasises the importance of using human *in vitro* systems to identify parasite effectors that mediate the pathogenesis of *Toxoplasma* in the human host. In both the RH and PRU screens we observed high levels of parasite restriction in HFFs, with ~90% of the mutant population restricted in cells pre-stimulated with 5U/ml IFNγ, a dose 20-fold less than that commonly used in literature. HFFs are extensively used for the continuous culture of *Toxoplasma* and are often considered as ‘non-immune’ cells, but these results support that structural cells can be important components of the innate immune response to *Toxoplasma* infection (Krausgruber *et al*., 2020). Other human non-hematopoietic cells have also been demonstrated to restrict *Toxoplasma* in response to IFNγ signalling, including neurons (Chandrasekaran *et al*., 2022), epithelial (Selleck *et al*., 2015) and endothelial cells (Clough *et al*., 2016).

High-content imaging of parasite restriction in HFFs revealed the RHΔMYR1 and RHΔMYR3 strains are highly susceptible to restriction through both increased vacuole clearance and growth restriction. Given the pleiotropic effects on the host that are mediated by MYR-dependent effector export (Naor *et al*., 2018), we assume that the abrogation of all effector export leads to hypersensitivity to multiple IFNγ-induced restriction pathways in HFFs. This is further supported as no other dense granule protein with an equally strong phenotype was identified in our screen, although it is possible that targets were filtered out due to variability between guides. As our library is constructed from hyperLOPIT prediction (Barylyuk *et al*., 2020), it could also be possible that some GRAs or ROPs are omitted if not previously annotated as localised to the dense granules or rhoptries. In contrast to MYR1 and MYR3 knockouts, single knockouts of GRA57, GRA70 or GRA71 were more sensitive to IFNγ-mediated vacuole clearance but did not display decreased replication in activated cells. As GRA57 was not shown to influence effector export, and both GRA57 and GRA71 do not appear to localise outside of the PV, we assume that this dense granule complex is not exported via the MYR translocon, and functions independently of the MYR complex to promote parasite survival of IFNγ.

In contrast to HFFs, we observed no significant impact on IFNγ survival in MEFs upon deletion of GRA57, with no previously identified phenotype for GRA57/GRA70/GRA71 in murine screens (Sangaré *et al*., 2019; Young *et al*., 2019; Y. Wang *et al*., 2020; Butterworth *et al*., 2022). From this we infer that the trimeric complex functions to subvert an IFNγ-induced response that is present in HFFs but absent or less dominant in MEFs. One such response is the induction of HFF cell death, leading to premature parasite egress (Niedelman *et al*., 2013). We observed a larger IFNγ-dependent decrease in vacuole numbers when HFFs were infected with knockouts of the GRA57/GRA70/GRA71 complex, an effect which could be explained by early egress of these strains in response to IFNγ. It has previously been shown that endolysosomal acidification of the parasitophorous vacuole mediates clearance of PRU parasites in IFNγ-activated HFFs, an effect that is inhibited through unknown mechanisms in RH parasites (Mukhopadhyay *et al*., 2020). Acidification of the PV during the normal lytic cycle of *Toxoplasma* serves as a signal for microneme secretion and parasite egress (Roiko, Svezhova and Carruthers, 2014). Therefore, acidification induced by host cell endolysosomal fusion with the PV may induce premature parasite egress, thereby limiting parasite replication and dissemination. Future work will therefore aim to determine if the GRA57/GRA70/GRA71 complex functions to resist IFNγ-induced vacuole acidification in HFFs.

Autophagy has emerged as a major clearance mechanism of *Toxoplasma* (Besteiro, 2019) and multiple other intracellular pathogens (Keller, Torres and Cadwell, 2020). In our efforts to determine if host autophagy contributed to the increased clearance of GRA57/GRA70/GRA71 knockouts, we unexpectedly found a marked reduction in the percent of these knockout vacuoles targeted by host ubiquitin in HFFs and MEFs, in both the presence and absence of IFNγ-stimulation. This phenotype does not provide a mechanistic explanation for the increased IFNγ-sensitivity upon knockout of GRA57/GRA70/GRA71, and further work is needed to clarify if there is a causal link between reduced ubiquitin recruitment and increased vacuole clearance of these strains.

Mukhopadhyay *et al*. have shown that ubiquitination of *Toxoplasma* vacuoles in HFFs is not intrinsically linked to downstream fusion with the endolysosomal system, therefore our observed decrease in vacuole ubiquitination may be a secondary effect of deletion of the GRA57/GRA70/GRA71 complex. However, one possible explanation for the observed decreased ubiquitination and increased parasite restriction is that GRA57/GRA70/GRA71 knockout strains become more rapidly targeted by endolysosomal acidification post-invasion, which could render partially disrupted vacuoles undetectable by standard immunofluorescence assays. At 3 hours post-infection, the total percentage of vacuoles with ubiquitin recruitment is reduced in these knockouts, but not entirely ablated (Figure 5C & 5D). As we stained for total mono- and poly-ubiquitin chain linkages, we do not yet know whether all vacuole ubiquitination is reduced simultaneously, or if specific linkages are depleted. Therefore, a further possibility is that in the absence of the complex, the vacuoles become hypersensitive to the remaining ubiquitination and subsequent disruption.

Recent work has found that RNF213, an IFNγ-induced host E3 ubiquitin ligase, mediates the majority of ubiquitin recruitment to both RH and PRU *Toxoplasma* vacuoles in HFFs (Hernandez *et al*., 2022; Matta *et al*., 2022); however, the target of this ligase is currently unknown. The observation that GRA57/GRA70/GRA71 deletion leads to reduced ubiquitination of the PV poses the intriguing question of whether the complex is required for RNF213 recruitment, is a direct target of ubiquitination, or is responsible for correct positioning of an RNF213 target at the PV. While each of the possibilities will be interesting to follow up to better understand the function of RNF213 in *Toxoplasma* restriction, they are likely not directly linked to the increased restriction of parasites upon GRA57 deletion, as reducing RNF213 activity by any means should increase, rather than decrease, *Toxoplasma* survival.

In conclusion, we identify several proteins that are required for survival in HFFs under conditions of IFNγ restriction in two *Toxoplasma* strains that differ in virulence. We identify a complex of three proteins that is required to protect *Toxoplasma* parasites from killing by the host cell. While deletion of the complex components leads to reduced ubiquitination of the parasite, we suggest that functional consequences of the complex deletion likely go beyond a role in ubiquitination.

It is interesting to note that none of the three proteins identified here to protect the parasite against the IFNγ response in human cells have been identified in previous screens in murine cells or mice. As humans are primarily an accidental host for *Toxoplasma*, it is highly likely that the GRA57 complex evolved to protect the parasite in species other than humans. It will be interesting to compare GRA57 and MYR function in cells of various origin to reveal commonalities in the cell autonomous immune response between humans and other species that can be infected by *Toxoplasma*.

## Methods

### CRISPR-Cas9 screens

#### Parasite transfections

To generate a pooled population of parasite effector knockouts, we used a plasmid library comprising 1299 single guide RNAs (sgRNAs) targeting 253 predicted secreted proteins, with an average of five sgRNAs/gene, with sgRNAs integrated into a pCas9-GFP-T2A-HXGPRT::sgRNA vector. The plasmid pool was linearised overnight with *Kpn*I-HF (NEB, R3142), purified by phenol-chloroform precipitation then resuspended in P3 (Lonza, V4XP-3024) transfection buffer at a concentration of 1ug/ul.

A minimum of 90 million parasites (RHΔHXGPRT or PRUΔHXGPRT) were transfected in triplicate using an Amaxa 4D Nucleofector (Lonza, AAF-1003X) with pulse code EO-115, with 30ug/transfection of the purified sgRNA plasmid library. Transfected populations were selected for after 24h using 25μg/mL mycophenolic acid (Sigma-Aldrich, 89287) and 50 μg/mL xanthine (Sigma-Aldrich, X3627) (M/X). Transfection efficiency was assessed via plaque assay, achieving a final guide coverage of 174X and 68X in the RH and PRU screens, respectively. Three days post-transfection, parasites were syringe lysed and added to HFF monolayers with 100 U/ml benzonase (Merck, E1014-25KU) overnight to remove traces of input DNA. Eight days post transfection, subset parasite populations were harvested for genomic DNA (gDNA) extraction to determine guide abundance in the starting inoculum. Remaining triplicate transfections were combined to generate the inoculum used for infections.

#### Infections

HFFs were grown to confluency in T175 flasks, then stimulated with 5U/ml human IFNγ (Bio-Techne, 285-IF-100) for 24h pre-infection. For infections, parasites were isolated from HFFs by syringe lysis through a 30-gauge needle (x3) and passed through 5μm filters (Millipore, SMWP04700), then added to HFFs at an MOI of 0.2 (1.4×10^6^ parasites/flask) for 48h. At 48h post-infection, parasites were harvested as above from HFFs, counted, then a subset were added to new flasks of IFNγ treatment-matched HFFs at an MOI of 0.2 for a further 48h. For each round of infection surviving parasites populations were expanded in unstimulated HFFs for a further 48h prior to harvesting for gDNA extraction.

#### gRNA isolation and sequencing

Genomic DNA was extracted from samples using Qiagen DNEasy Blood kit, then guide sequences were amplified from gDNA and the plasmid pool by nested PCR using KAPA HIFI Hotstart PCR kit (Kapa Biosystems, KK2501), as previously described in (Young *et al*., 2019). Primers used for nested PCR are listed in Supplementary Data 4 (primers 1-20). Purified PCR products were then sequenced on a HiSeq400 (Illumina) with single end 100 bp reads at a minimum read depth of 5 million reads/sample.

#### Sequencing data analysis

gRNA sequences were aligned to a reference library as described previously in (Young *et al*., 2019; Butterworth *et al*., 2022). The lowest 1.5 percentile of guides expressed across all samples were removed from the analysis. Counts were normalised using the median of ratios, then genes represented by fewer than 3 matching guides were removed from the analysis. The median L2FC for each gene was calculated from the normalised counts at the end of growth in pre-stimulated HFFs relative to unstimulated HFFs, or from counts at the end of growth in unstimulated HFFs relative to the counts in the starting inoculum sample. The median absolute deviation (MAD) score across gRNA L2FCs targeting each gene was calculated, and genes with the highest 1.5% of MAD scores were removed from the analysis. A DISCO score based on the local FDR-corrected q-value was calculated for each L2FC to compare between untreated and IFNγ pre-stimulated HFFs.

### Parasite and host cell culture

*Toxoplasma gondii* strains were maintained by serial passage every 2-3 days in HFFs (ATCC, SCRC-1041). For experiments parasites were isolated by syringe lysis through a 27-gauge needle and passed through 5μm filters. Parasite genotypes were verified by restriction fragment length polymorphism analysis of the SAG3 gene using primers 70 and 71 (Su, Zhang and Dubey, 2006). HFFs and MEFs (gift from Felix Randow) were maintained in DMEM with GlutaMAX (Gibco) supplemented with 10% foetal bovine serum (Gibco). Parental strains used were RHΔHXGPRT and PRUΔHXGPRT (Donald *et al*., 1996), RHΔKU80 (Huynh and Carruthers, 2009), and PRUΔKU80 (Fox *et al*., 2011).

### Generation of parasite cell lines

#### Knockouts

For generation of RHΔGRA57 and PRUΔGRA57 strains, two guides were designed targeting exon 1 and exon 7 of the GRA57 coding sequence (CDS). Guides were integrated separately into a pCas9-GFP::sgRNA vector by inverse PCR using a general reverse primer (21) and primers 24 or 25. The exon 1 targeting plasmid was digested with XhoI/KpnI (NEB), and the exon 7 guide was amplified with primers 22 and 23 to facilitate gibson cloning of the exon 7 guide into the same backbone. For all other knockouts generated in this study (RHΔMYR3, RHΔGRA70 and RHΔGRA71), single guide plasmids were generated using primers 26-28. Homology repair cassettes were generated by PCR amplification from Pro^GRA1^-mCherry-T2A-HXGPRT-Ter^GRA2^, using primers with 40bp flanking homology to the 5’ and 3’ UTRs of each gene (primers 29-36). 10ug of repair cassette was purified with 5ug of pCas9-GFP::sgRNA and transfected into RHΔKu80 or PRUΔKu80 parasites, using an Amaxa 4D Nucleofector as described above. Transfectants were selected 24h post-transfection with 25μg/ml of mycophenolic acid (Merck) and 50μg/ml xanthine (Sigma) (M/X) and cloned by limiting dilution. Integration of the mCherry-HXGPRT repair templates was verified by PCR using primers 37-46.

#### Complementation

To complement the RHΔGRA57 strain, the 5’UTR and CDS of GRA57 was generated through a combination of PCR amplification from gDNA (5’UTR, exons 1 and 7-primers 47-54), and amplification from gBlocks (IDT) for exons 2-4 and exon 6 (listed in Supplementary Data 4). PCR products were inserted into the pUPRT vector by Gibson assembly. The pUPRT:GRA57-HA plasmid was linearised with ScaI, then 10ug was transfected into RHΔGRA57 alongside 1ug of pCas9-GFP::UPRT. Transfectants were selected 24h post-transfection with 5μM 5’-fluo-2’-deoxyuridine (FUDR), then cloned and verified by PCR at the UPRT locus using primers 55-58.

#### Endogenous tagging

To endogenously tag GRA57 with an HA epitope tag at the C-terminus, a pCas9-GFP::sgRNA plasmid targeting the 3’end of exon 7 of GRA57 was generated by inverse PCR from pCas9-GFP::sgRNA using primers 21 and 59. A repair cassette with 40bp flanking homology was amplified by PCR from HA-Ter^GRA2^::Pro^DHFR^-HXGPRT-Ter^DHFR^ using primers 61 and 62. 10ug of repair cassette was purified with 5ug of pCas9-GFP::sgRNA and transfected into RHΔKu80 as above. For endogenous tagging of GRA70 with a C-terminal V5 tag, the pCas9-GFP::sgRNA plasmid was generated using primers 21 and 60, and the repair cassette was generated by PCR from a V5-Ter^GRA2^::Pro^DHFR^-HXGPRT-Ter^DHFR^ construct, using primers 63 and 64. Transfectants were selected 24h post-transfection with 25μg/ml of mycophenolic acid (Merck) and 50μg/ml xanthine (Sigma) (M/X), then cloned and verified by PCR using primers 65-69.

All primers used for generating parasite lines are listed in Supplementary Data 4

### Plaque assays

100 parasites were added to HFF monolayers in a T25 flask and allowed to grow undisturbed for 10 days. Cells were fixed with 100% methanol and stained with crystal violet, then plaques were imaged on a ChemiDoc imaging system (BioRad). Plaque area was measured in FIJI (Schindelin *et al*., 2012). Differences between strains were determined by one-way analysis of variance (ANOVA) with Tukey’s multiple comparison test.

### Co-immunoprecipitation mass-spectrometry

#### Immunoprecipitation

HFFs grown to confluency in T175 flasks were pre-stimulated with 2.5U/ml IFNγ (Bio-Techne, 285-IF-100) for 6 hours prior to infection with RHΔKu80 or RHGRA57-HA in triplicate. 24h post-infection, infected cells were washed 3x in cold PBS then lysed in cold immunoprecipitation (IP) buffer (10mM Tris, 150mM NaCL, 0.5mM EDTA + 0.4% NP40, pH 7.5 in H_2_O, supplemented with 2x cOmplete Mini EDTA-free Protease Inhibitor Cocktail). Lysates were syringe lysed 6x through a 30g needle, then centrifuged at 2000g for 20 minutes to remove the insoluble fraction. Soluble fractions were added to 50ul/sample anti-HA agarose matrix (Thermo), then incubated overnight at 4°C with rotation. The matrix was washed three times with IP buffer, then proteins were eluted in 30μL 3x Sample Loading Buffer (NEB) at room temperature for 10 minutes.

#### Mass spectrometry

20ul of each IP elution was loaded on a 10% Bis-Tris gel and run into the gel for 1cm, then stained with InstantBlue Coomassie Protein Stain. Proteins were alkylated in-gel prior to digestion with 100ng trypsin (modified sequencing grade, Promega) overnight at 37°C. Supernatants were dried in a vacuum centrifuge and resuspended in 0.1% TriFluoroAcetic acid (TFA). 1-10ul of acidified protein digest was loaded onto a 20mm x 75um Pepmap C18 trap column (Thermo Scientific) on an Ultimate 3000 nanoRSLC HPLC (Thermo Scientific) prior to elution via a 50cm x 75um EasySpray C18 column into a Lumos Tribrid Orbitrap mass spectrometer (Thermo Scientific). A 70’ gradient of 6%-40%B was used to elute bound peptides followed by washing and re-equilibration (A= 0.1% formic acid, 5% DMSO; B= 80%ACN, 5% DMSO, 0.1% formic acid). The Orbitrap was operated in “Data Dependent Acquisition” mode followed by MS/MS in “TopS” mode using the vendor supplied “universal method” with default parameters.

Raw files were processed to identify tryptic peptides using Maxquant (maxquant.org) and searched against the *Toxoplasma* (ToxoDB-56_TgondiiGT1_AnnotatedProteins) and Human (Uniprot, UP000005640) reference proteome databases and a common contaminants database. A decoy database of reversed sequences was used to filter false positives, at peptide and protein false detection rates (FDR) of 1%. T-test based volcano plots of fold changes were generated in Perseus (maxquant.net/perseus) with significantly different changes in protein abundance determined by a permutation based FDR of 0.05% to address multiple hypothesis testing.

### Protein secondary structure prediction

Protein sequences for GRA57, GRA70 and GRA71 from ToxoDB were used to predict boundaries and regions of secondary structure with PSIPRED (Buchan and Jones, 2019). Transmembrane domains (TMDs) were assigned using MEMSAT-SVM (within PSIPRED), DeepTMHMM (J *et al*., 2022) and CCTOP (Dobson, Reményi and Tusnády, 2015), with TMDs only annotated where ≥2 programmes concurred. Regions of predicted coiled-coil were determined using Waggawagga (Simm, Hatje and Kollmar, 2015), with coiled-coil annotated where the probability score was ≥ 90.

### Western blotting

Parasites were isolated from host cells by syringe-lysis through 27-gauge needles, passed through 5μm filters and washed x1 with cold PBS. Samples were lysed on ice in 1% NP40 IP buffer (10mM Tris, 150mM NaCL, 0.5mM EDTA + 1% NP40, pH 7.5 in H_2_O, supplemented with 1x cOmplete Mini EDTA-free Protease Inhibitor Cocktail) for 30 minutes, then spun at 2000g for 15 minutes to remove the insoluble fraction. 10ug of protein per sample was incubated with 3x loading buffer for 10 minutes at room temperature, then separated by SDS-PAGE on a NuPAGE^™^ 3 to 8%, Tris-Acetate gel. Proteins were transferred to a nitrocellulose membrane using the High Molecular Weight protocol on the Trans-Blot Turbo transfer system (Bio-Rad). Membranes were blocked in 5% milk in 0.05% Tween 20 in PBS for 1h at room temperature, followed by incubation for 1h at room temperature with primary antibodies diluted in 1% milk in 0.05% Tween 20 in PBS. Blots were then incubated with appropriate secondary antibodies for 1h at room temperature. Primary antibodies used were 1:200 mouse anti-T. gondii (Santa Cruz, SC-52255), 1:1000 rat anti-HA-Peroxidase (Roche, 12013819001) and 1:1000 rabbit anti-V5 (Abcam, AB_2809347). Secondary antibodies used were 1:10000 goat anti-mouse HRP (Insight Biotechnologies, 474-1806) and 1:3000 goat anti-rabbit HRP (Insight Biotechnology, 474-1506). HRP was detected using an enhanced chemiluminescence (ECL) kit (Pierce), visualised on a ChemiDoc imaging system (BioRad).

### Transmission electron microscopy

HFFs were grown to confluency on 13mm glass coverslips. HFFs were infected with 60,000 parasites per strain for 24h, then washed 1x with DPBS prior to fixation in 2.5% glutaraldehyde + 4% formaldehyde in 0.1M phosphate buffer (PB) for 30 minutes. Samples were washed 2x with 0.1M PB and stained with 1% (v/v) osmium tetroxide (Taab) / 1.5% (v/v) potassium ferricyanide (Sigma) for 1 hour at 4°C. Samples were washed 2x in dH2O and were then transferred to a Pelco BioWave Pro+ microwave (Ted Pella Inc, Redding, USA) for a further two dH2O washes in the Biowave without vacuum (at 250W for 40s). The SteadyTemp plate was set to 21°C. In brief, the samples were incubated in 1% (w/v) tannic acid in 0.05M PB pH 7.4 (Sigma) for 14 minutes under vacuum in 2 minute cycles alternating with/without 100W power, followed by 1% sodium sulphate in 0.05M PB pH 7.4 (Sigma) for 1 minute without vacuum at 100W. The samples were then washed in dH2O and dehydrated in a graded ethanol series (25%, 50%, 70%, 90%, and 100%, twice each), and in acetone (3 times) at 250 W for 40s without vacuum. Exchange into Epon 812 resin (Taab) was performed in 25% 50% and 75% resin in acetone, at 250 W for 3 minutes, with vacuum cycling (on/off at 30 sec intervals). The samples were transferred to 100% resin overnight before embedding at 60°C for 48 h. 70 nm sections were sliced with a diamond knife on a RMC Powertome Ultramicrotome, and sections analysed by a JEM-1400 FLASH transmission electron microscope (Jeol) with Jeol Matataki Flash sCMOS camera.

### Immunofluorescence assays

HFFs were seeded to confluency on 15mm glass coverslips and infected with 60,000 parasites/well for 24h. Cells were fixed in 4% paraformaldehyde (PFA) for 15 minutes, then permeabilised in 0.2% Triton-X100 (GRA2 staining) or in 0.1% saponin (GRA3 staining) for 10 minutes and blocked with 3% bovine serum albumin (BSA) in DPBS for 30 minutes. Primary and secondary antibodies used for immunofluorescence assays are listed in Supplementary Data 6. All antibodies were incubated in 3% BSA in DPBS for 1h at room temperature, with 3x washes in DPBS between stages. Final secondary staining was combined with DAPI (5ug/ml). Slides were mounted in ProLong Gold (ThermoFisher, P36930). Images were acquired on a VisiTech instant SIM (VT-iSIM) microscope using a 150X oil-immersion objective with 1.5μm z axis steps. Resultant stacks were deconvoluted and processed using Microvolution plugin in FIJI (Schindelin *et al*., 2012).

### c-Myc nuclear translocation assays

HFFs were grown to confluence in 8-well ibidi μ-slides (Ibidi, 80806). Cells were infected with 30,000 parasites in DMEM with 0.1% FBS. After 24h, slides were fixed in 4% paraformaldehyde (PFA) for 15 minutes, permeabilised in 0.2% Triton-X100 for 10 minutes then blocked in 3% BSA for 30 minutes. Host c-Myc was stained using 1:800 rabbit anti-cMyc for 2h at room temperature, followed by 1:1000 anti-Rabbit-AlexaFluor488 and 5μg/mL DAPI for 1h at room temperature. Images were acquired on a Nikon Ti-E inverted widefield fluorescence microscope with a Nikon Plan APO 40x/0.95 objective and Hamamatsu C11440 ORCA Flash 4.0 camera. Host nuclear c-Myc signal of infected cells was measured in FIJI, and the median background c-Myc signal was subtracted for each image. Data is shown as the median fluorescence intensity for each strain relative to RHΔUPRT in each biological replicate. Differences between strains were determined by one-way analysis of variance (ANOVA) with Tukey’s multiple comparison test.

### Cytation plate reader assays

#### Survival assays

Host cells were seeded in 96-well black imaging plates (Falcon) to confluency, then media was changed to phenol red free DMEM (Gibco, A1443001) with or without 100U/ml IFNγ (Human:Bio-Techne, 285-IF-100| Mouse: ThermoFisher, Gibco PMC4031) for 24 hours. Parasite strains were syringe lysed and added to host cells at 40,000 parasites/well. Plates were imaged live at 24h post-infection (MEFs) or 48h post-infection (HFFs) on a Cytation 5 plate reader using the 4X objective (HFFs) or 20X (MEFs). Total area with signal in the Texas Red channel (Ex/Em 586/647) was measured per well. Differences between strains were determined by paired two-sided t-test.

#### Tryptophan supplementation

HFF were seeded as described above, then treated wells were supplemented with 1mM L-tryptophan (VWR, J62508-09) dissolved in 0.1N NaOH at the same time as IFNγ stimulation, as described in (Niedelman *et al*., 2013). Untreated wells had 0.1N NaOH added as a vehicle control. Data acquisition on a Cytation 5 plate reader was performed as with survival assays.

### High-content imaging survival assays

Host cells were seeded to confluency in 96-well imaging plates (Ibidi). Cells were pre-stimulated for 24h with 100U/ml IFNγ according to host species. Cells were infected at an MOI of 0.3 for 24h, then fixed in 4% PFA and stained with 5ug/ml DAPI and 5ug/ml CellMask Deep Red (Invitrogen). Plates were imaged using the Opera Phenix high-content screening system, with 25 images and 5 focal planes acquired per well. Automated analysis of infection phenotypes was performed using Harmony v5 (PerkinElmer) as described in (Butterworth *et al*., 2022). Data is reported as the mean proportion of each factor (total *Toxoplasma* number, vacuole size or vacuole number) in IFNγ treated wells relative to untreated wells. Differences between strains were determined by paired two-sided t-test.

### Vacuole ubiquitination assay

Host cells were seeded and pre-stimulated as above in 96-well imaging plates (Ibidi). Cells were infected with 80,000 parasites/well and centrifuged at 300g for 5 minutes to synchronise infection. At 3 hours post-infection, cells were washed 3x in DPBS to remove extracellular parasites, then fixed in 4% PFA for 15 minutes and blocked with 3% BSA in DPBS for 1 hour at room temperature. Extracellular parasites were stained prior to permeabilisation using 1:1000 rabbit anti-toxo (Abcam, ab138698) and 1:500 goat anti-rabbit-AlexaFluor647 (Life Technologies, A21244). Cells were then permeabilised with 0.2% Triton-X100 for 10 minutes and re-blocked with 3% BSA. Total ubiquitin was stained using 1:200 mouse anti-ubiquitin (Merck, ST1200) for 2 hours at room temperature, followed by donkey anti-mouse-AlexaFluor488 (Thermo, A32766) for 1 hour at room temperature. Plates were imaged using the Opera Phenix high-content screening system, with 25 images and 5 focal planes from −1 to 1 μm with a step size of 0.5 μm acquired per well. Automated analysis was performed using Harmony v5 (PerkinElmer). Recruitment was automatically quantified by first excluding extracellular parasites, then measuring AF488 signal within the vacuole region (1.5μm boundary of vacuole) and the cytoplasmic region around the vacuole (2 to 3μm boundary of vacuole). Parasites with AF488 signal 1.3X higher within this radius relative to the cytoplasmic background of each infected cell were classed as ubiquitin-recruited. Image acquisition parameters and analysis sequence are detailed further in Supplementary Data 5. For each well the percentage of ubiquitin-recruited intracellular vacuoles was determined, then the mean percentage recruitment was calculated across triplicate wells. Differences between strains were determined by paired two-sided t-test.

### RNA-Seq

#### RNA sample preparation

HFFs in T25 flasks were serum starved for 24h with DMEM containing 0.5% FBS. Treated flasks were simultaneously pre-stimulated for 24h with 100U/ml IFNγ (Bio-Techne, 285-IF-100). For infections, parasites were isolated by syringe lysis through a 30-gauge needle, passed through 5μm filters, centrifuged and resuspended in 0.5% FBS DMEM. HFFs were infected in triplicate with 1 million parasites per flask. 24 hours post-infection, samples were collected by washing each flask x1 in cold PBS, scraping in 2ml PBS and transferring to an RNAse free tube. Samples were centrifuged at 2000rpm for 10 minutes, then lysed in 600ul RLT buffer. Lysates were homogenised using Qiashredders (Qiagen, 79656), then RNA was isolated using an RNeasy Mini Kit (Qiagen, 74104) according to manufacturer’s instructions. mRNA libraries were prepared using the NEBNext Ultra II Directional PolyA mRNA kit (NEB, E7760L) with 100ng of input, then sequenced on a NovaSeq (Illumina) using paired end 100bp reads to a minimum depth of 30 million reads per sample.

#### RNA-Seq data analysis

RNA-seq data was quantified using STAR/RSEM (Li and Dewey, 2011) from within the nfcore/rnaseq (Ewels *et al*., 2020) pipeline (version 3.4), against human and *Toxoplasma* transcriptomes (GRCh38, annotation release-95 from Ensembl and ToxoDB-59_TgondiiGT1 obtained from ToxoDB). Differential analysis was run across strain and IFNγ treatment groups using DESeq2 (1.36.0) (Love, Huber and Anders, 2014), correcting for experimental batch effect in the model. Pairwise comparisons were run, and an interactions analysis across the two experimental factors against uninfected samples and -IFNγ control group in each case. RSEM counts were imported using tximeta (1.14.1) to account for transcript length, and IHW (1.24.0) was used to control from multiple testing in the differential gene selection (< 0.05 FDR). Shrunken log fold changes were calculated using type=“ashr”.

## Supplementary material

### Supplementary figures

**Figure S1:**
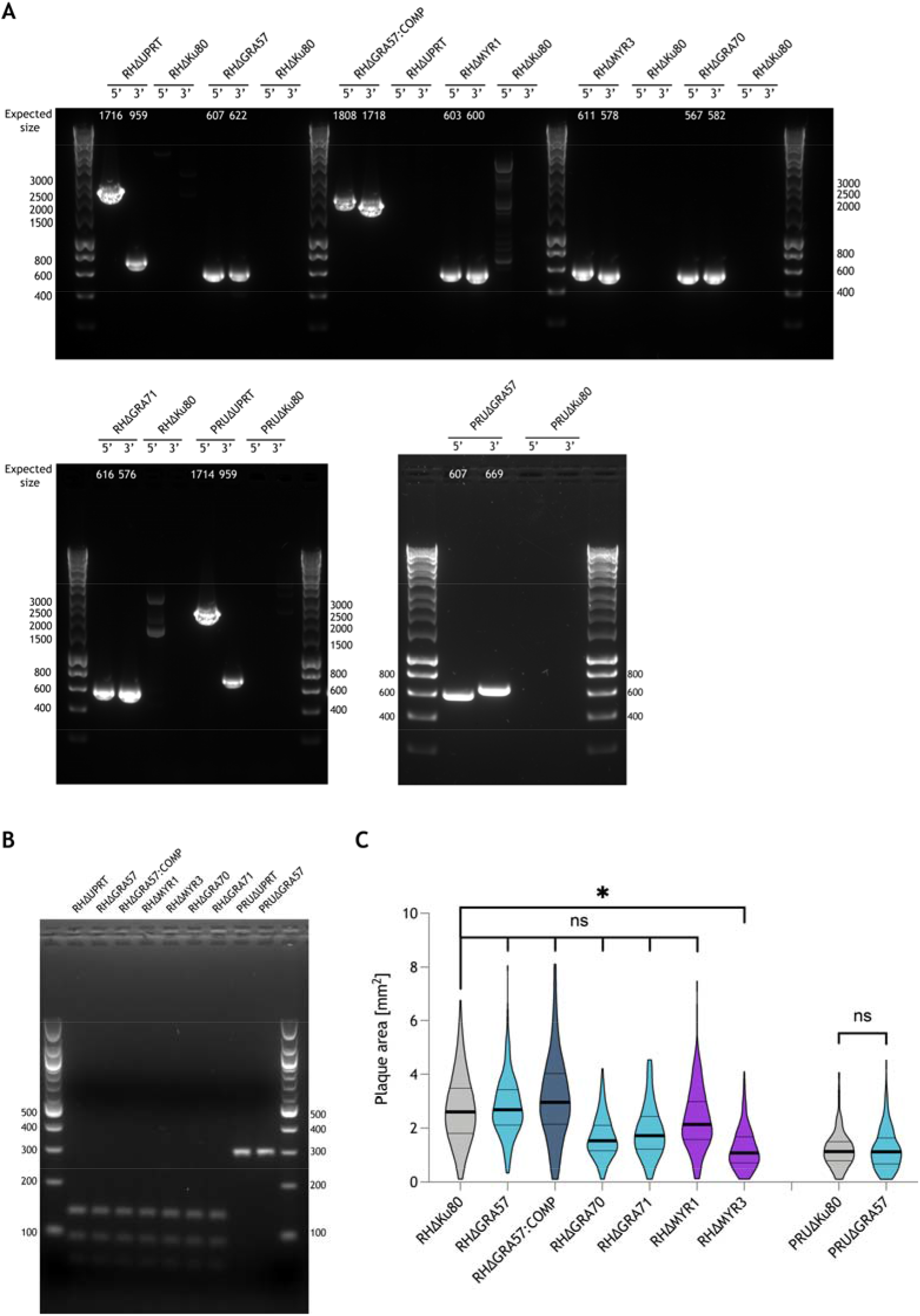
Verification of *Toxoplasma* knockout lines generated in this study. **(A)** PCR verification of successful integration of repair cassettes at indicated gene loci. Primers used for verification are listed in supplementary table 4. **(B)** Strain genotype verification by restriction fragment length polymorphism (RFLP) of the SAG3 gene **(C)** Quantification of plaque area after 10 days growth in HFFs. Results are shown as violin plots with median and quartiles from a minimum of 3 biological replicates. p-values were calculated by one-way analysis of variance (ANOVA) with Tukey’s multiple comparison test. *, p< 0.05; ns, not significant.

**Figure S2:**
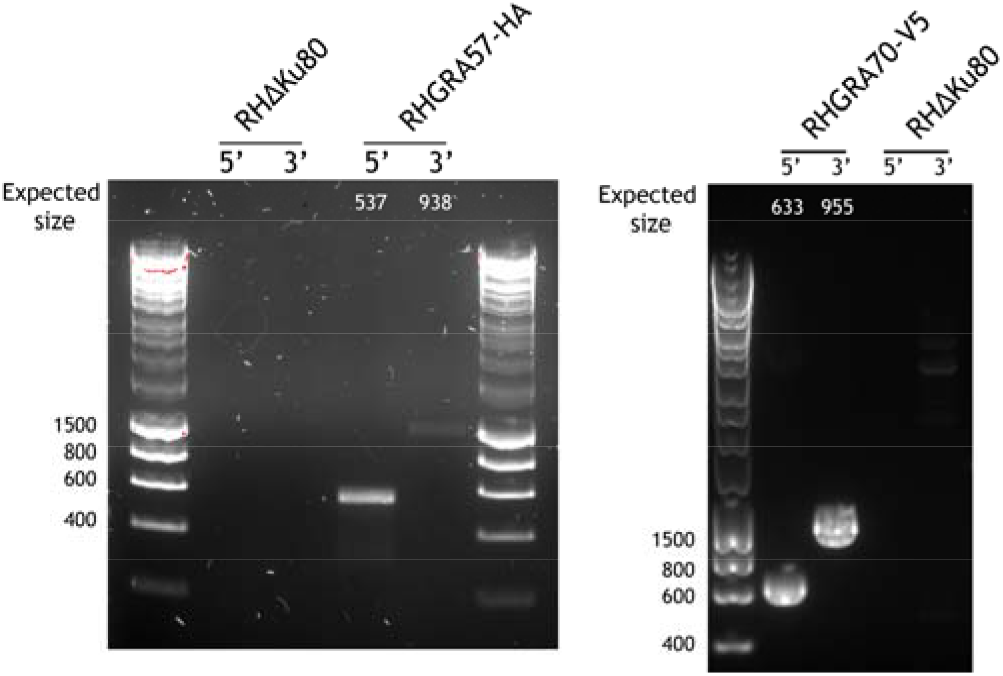
PCR verification of endogenously c-terminally tagged *Toxoplasma* lines generated in this study. Primers used for verification are listed in supplementary table 4.

**Figure S3:**
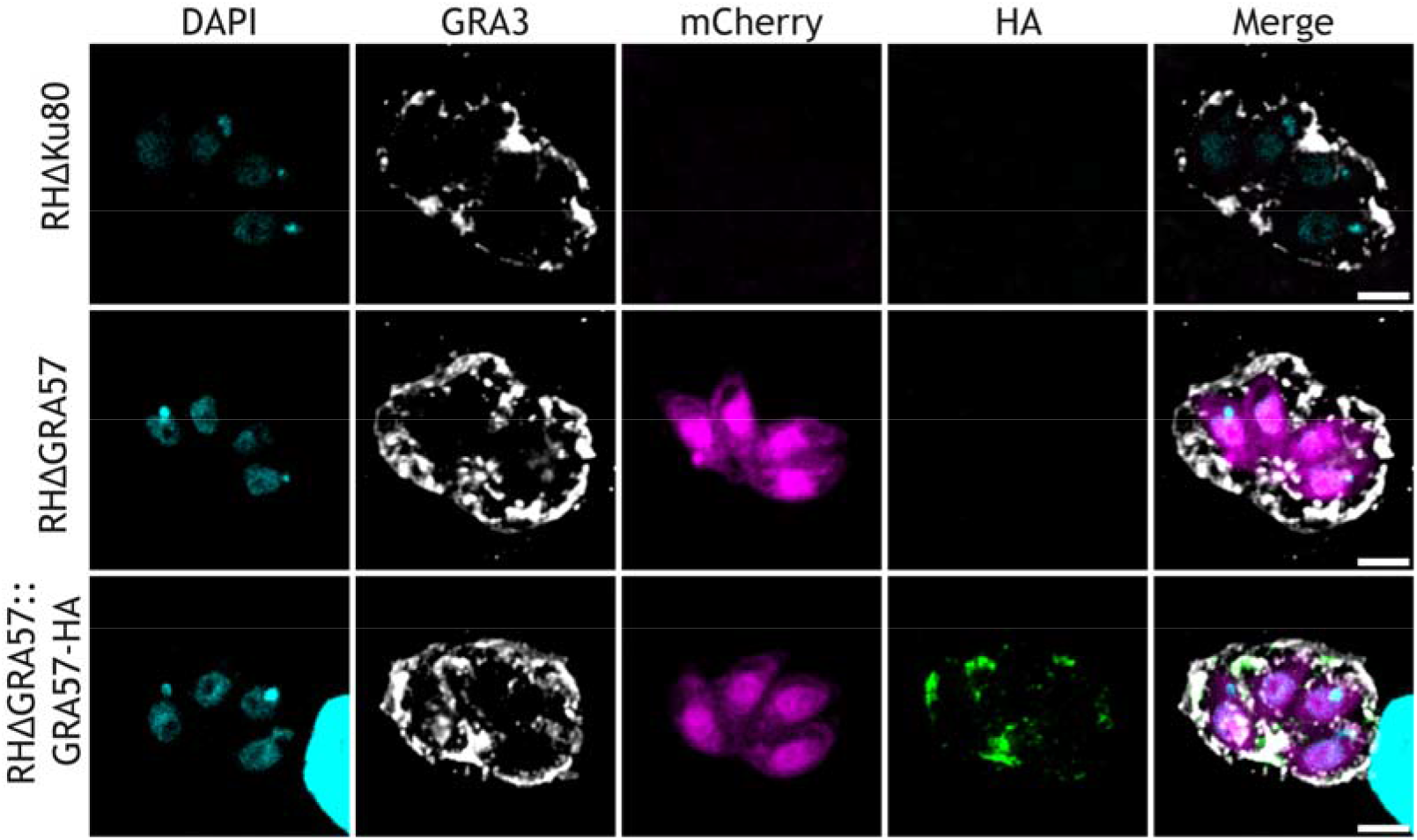
GRA57-HA partially co-localises with the PVM marker GRA3. Scale bar represents 3um.

**Figure S4:**
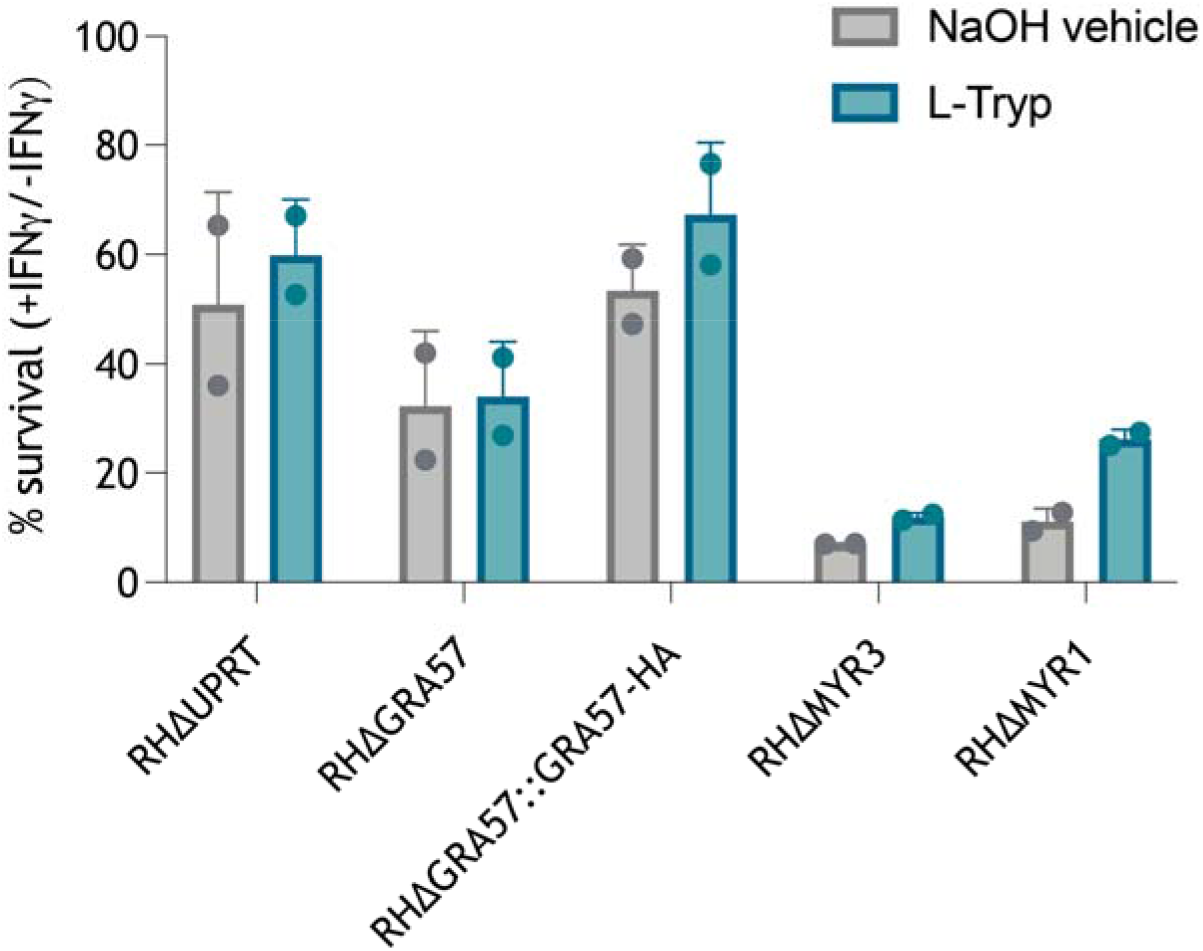
Survival of ΔGRA57 parasites is not rescued with exogenous supplementation of tryptophan. HFF IFNγ restriction assays as in Figure 2, with the addition of 1mM L-tryptophan to treated conditions simultaneous with IFNγ pre-stimulation. Untreated controls had 0.1 N NaOH added as a vehicle control. Host cells were infected in technical triplicate with the indicated parasite strains for 24h at an MOI of 0.3, then imaged live on a Cytation 5 plate reader. Total mCherry signal area per well was measured to determine parasite growth in IFNγ stimulated relative to unstimulated cells. Data displayed as mean survival + standard deviation from two biological replicates.

**Figure S5:**
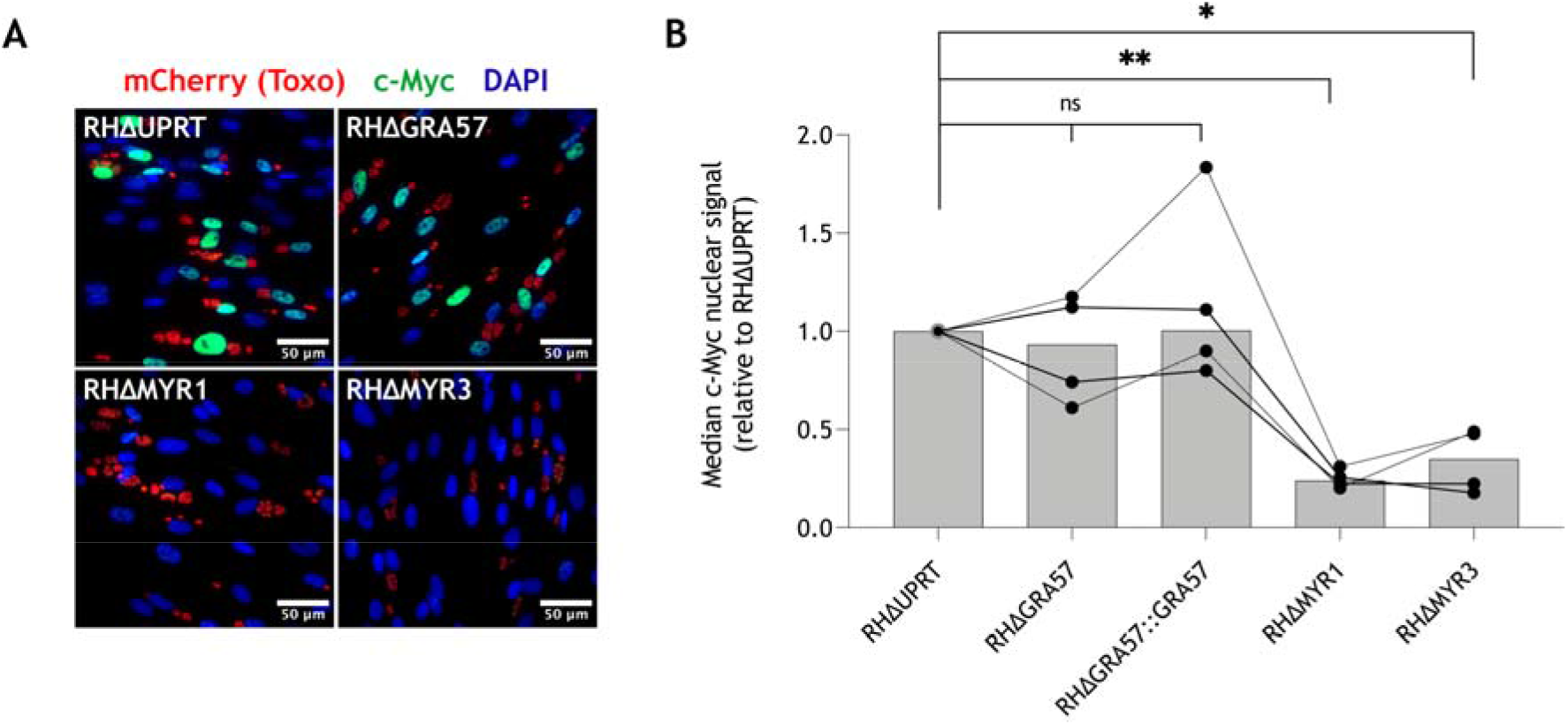
GRA57 does not contribute to MYR-dependent translocation of host c-Myc. **(A)** Representative IFA images. HFFs were infected for 24h prior to fixation and staining for host c-Myc. **(B)** Quantification of nuclear translocation of host c-Myc. Median nuclear c-Myc signal in infected cells was measured in FIJI, with the median background c-Myc signal subtracted for each image. Median nuclear c-Myc fluorescence intensity for each strain was normalised to that of RHΔUPRT in each biological replicate. Data is shown as median with individual biological replicates overlayed. p-values were calculated by one-way analysis of variance (ANOVA) with Tukey’s multiple comparison test. *, p< 0.05; **, p < 0.01; ns, not significant.

**Figure S6:**
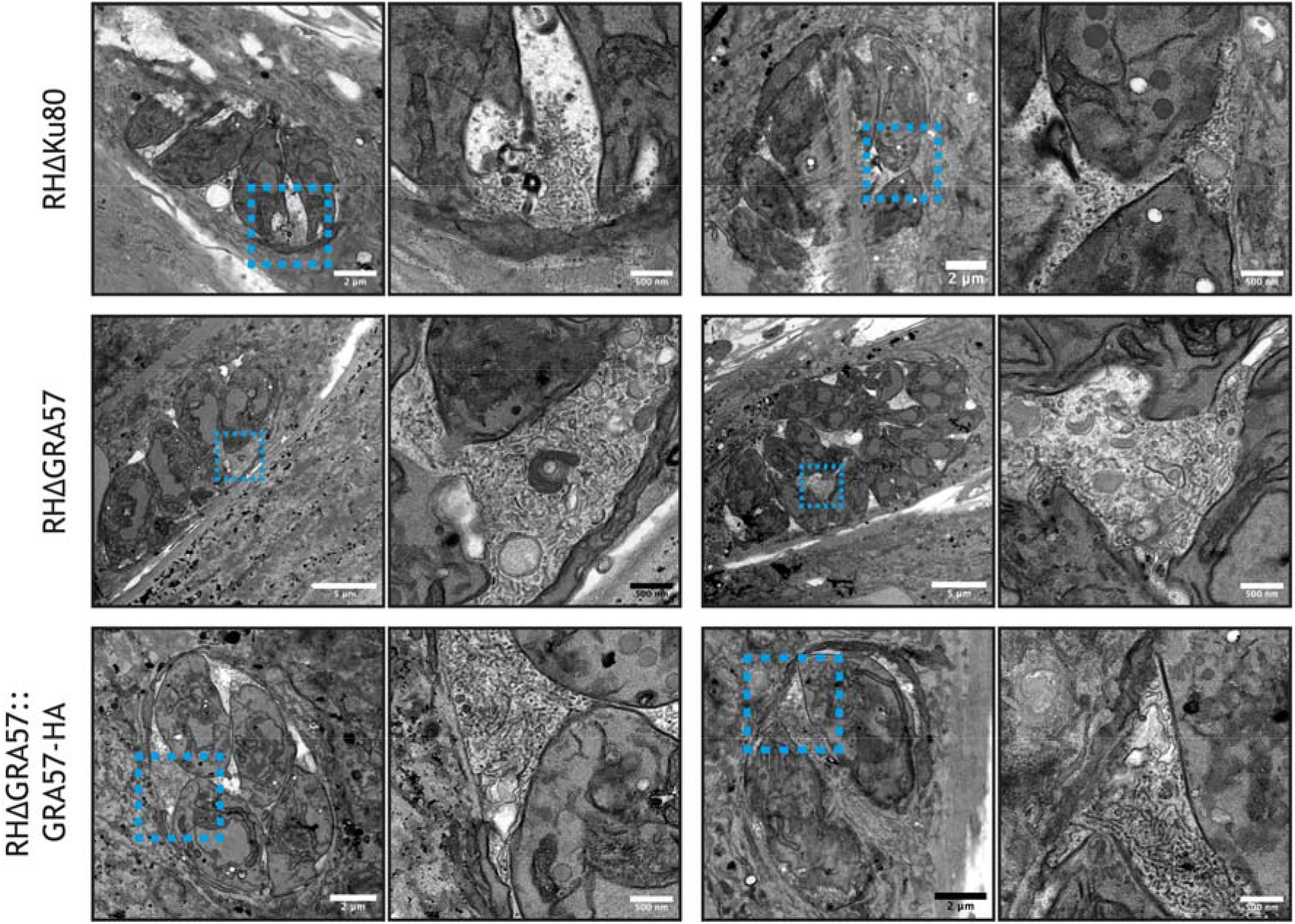
GRA57 deletion does not affect formation of the IVN. HFFs monolayers were infected with the indicated strains for 24h prior to fixation and preparation for transmission electron microscopy (TEM).

**Figure S7:**
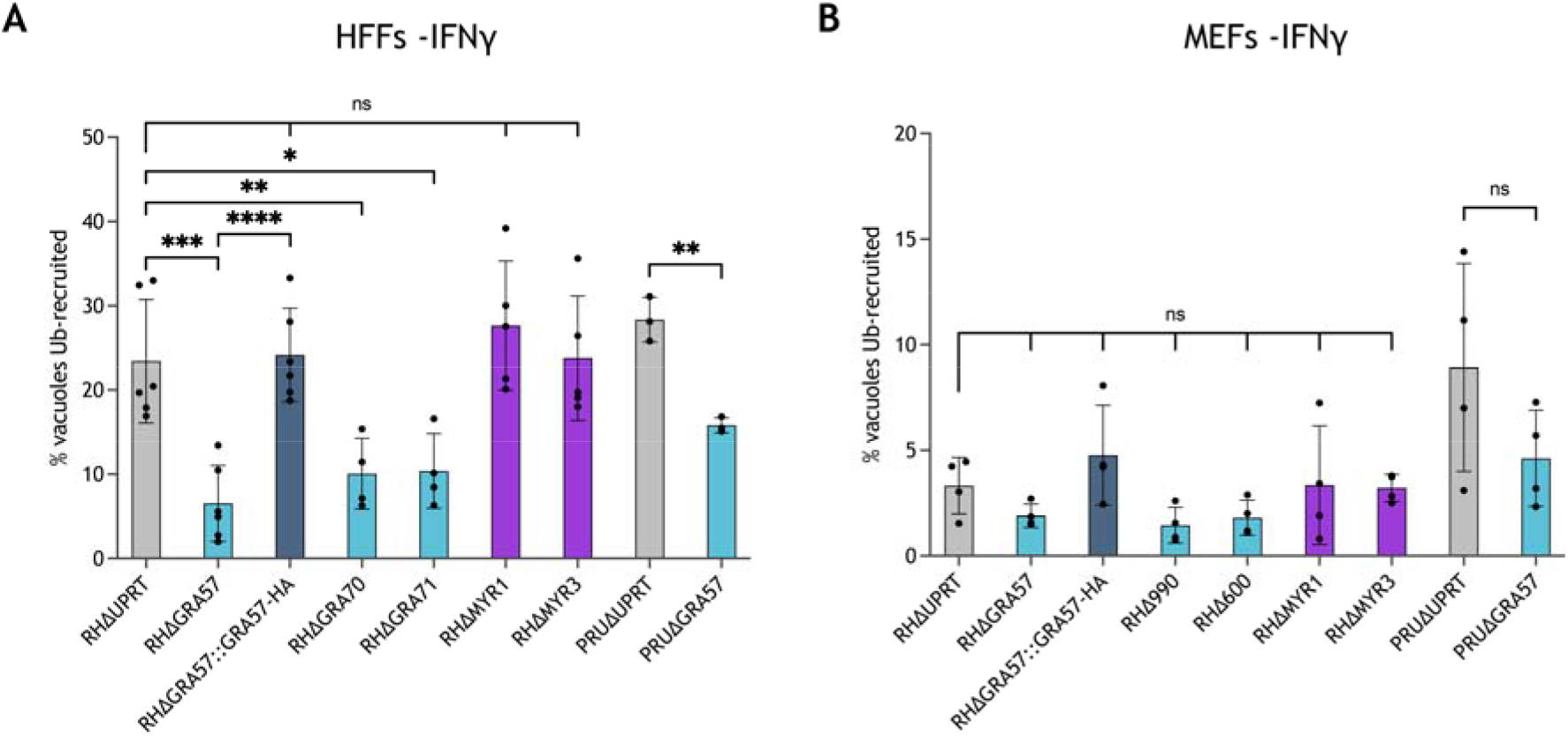
Deletion of GRA57, GRA70 or GRA71 leads to reduced host ubiquitin recruitment in unstimulated cells. Recruitment of host ubiquitin to *Toxoplasma* vacuoles in (**A)** HFFs and (**B)** MEFs. Related to Figure 5, data shows ubiquitin recruitment levels in unstimulated cells. Recruitment of ubiquitin was automatically counted using high content imaging and analysis. p-values were calculated by paired two-sided t-test. *, p < 0.05; **, p < 0.01; ***, p < 0.001; ****, p < 0.0001; ns, not significant.

**Figure S8:**
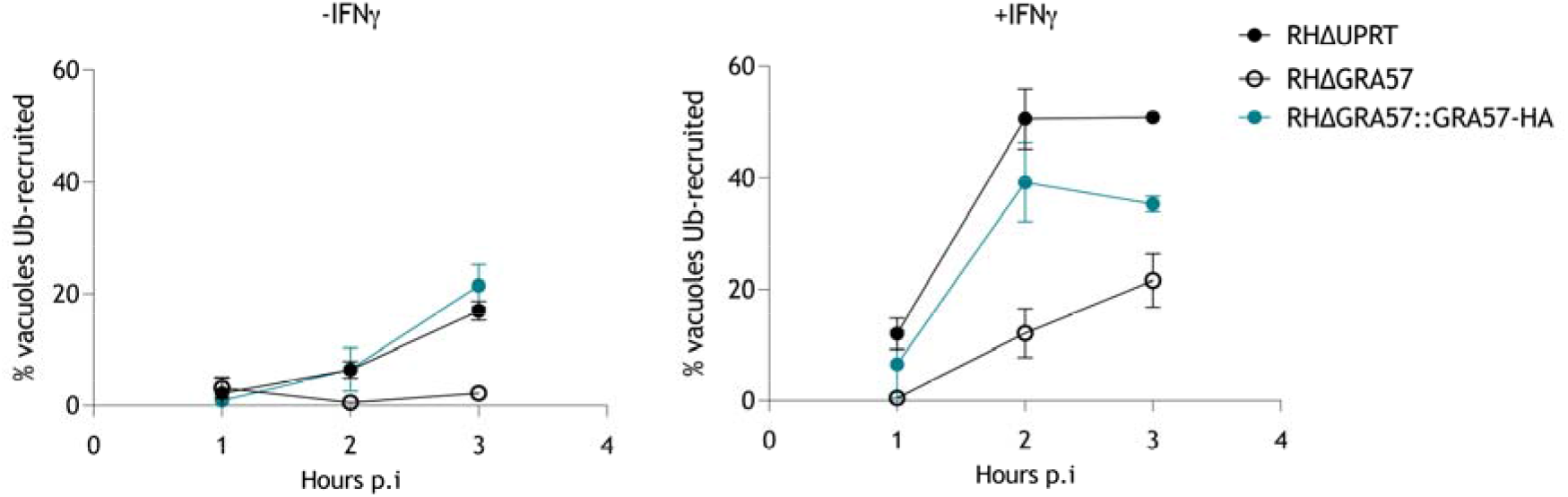
RHΔGRA57 vacuoles are targeted less frequently by host ubiquitin during the early stages of infection. HFFs were infected as in Figure 5, then fixed at the indicated time points postinfection. Data shown as mean of technical triplicate ± standard deviation.

## Supplementary Data

**Supplementary Data 1:** Data from CRISPR screens in RH and PRU parasites. Raw read counts, normalised counts, guide L2FCs, gene L2FCs, p-values, and DISCO scores.

**Supplementary Data 2:** Co-immunoprecipitation and mass spectrometry results.

**Supplementary Data 3:** RNASeq of infected HFFs. Differential gene analysis and interaction analysis. IFNγ treatment is denoted as yes or no. For the interaction analysis, sheet names represent the IFNγ effect for the first strain relative to the IFNγ effect for the second strain (ie WT|IFN_vs_dGRA57|IFN represents +IFNγ vs - IFNγ in the RHΔKu80 group relative to +IFNγ vs -IFNγ in the RHΔGRA57 group.

**Supplementary Data 4:** Primer sequences used in this work.

**Supplementary Data 5:** Opera Phenix image acquisition parameters and Harmony analysis sequence used for automated ubiquitin recruitment analysis.

**Supplementary Data 5:** Antibodies used for immunofluoresence assays.

## Acknowledgements

We thank all members of the Treeck laboratory as well as Barbara Clough and Eva Frickel for critical discussions, and Stephanie Nofal for critically reading the manuscript. We thank Michael Howell, Rachael Instrell and Becky Saunders (High-Throughput Screening Science Technology Platform, The Francis Crick Institute, London, United Kingdom) for assistance with sgRNA library preparation. We thank members of the Advanced Sequencing, High Throughput Screening, Electron Microscopy, Proteomics and Cell Services Science Technology Platforms at the Francis Crick Institute for support. We thank Bishara Marzook for assistance with the Cytation 5 plate imager, Jean Francois Dubremetz for providing the GRA3 antibody, Felix Randow and Ana Crespillo-Casado for providing the MEFs and Matthew Cottee for assistance with protein structural analysis. We acknowledge ToxoDB (http://toxodb.org/) for providing an invaluable resource that made this work possible.

This work was supported by funding to MT from the Francis Crick Institute which receives its core funding from Cancer Research UK (FC001189), the UK Medical Research Council (FC001189), and the Wellcome Trust (FC001189). The Science Technology Platforms at the Francis Crick Institute receive funding from Cancer Research UK (FC001999), The UK Medical Research Council (FC001999) and the Wellcome Trust (FC001999). For the purpose of Open Access, the authors have applied a CC BY public copyright licence to any Author Accepted Manuscript version arising from this submission.

## Author contributions

Conceptualisation: EJL, FT, MT. Investigation: EJL, FT, SB, OKS, SH, AW. Formal analysis: EJL, FT, SB, OKS, SH, PE, MT. Visualisation: EJL Supervision: MT. Project administration: MT. Funding acquisition: MT. Writing - original draft: EJL, MT. Writing - review and editing: all authors.

